# A co-speciation dilemma and a lifestyle transition with genomic consequences in *Wolbachia* of Neotropical *Drosophila*

**DOI:** 10.1101/2025.04.16.649125

**Authors:** Konstantinos Papachristos, Wolfgang J. Miller, Lisa Klasson

## Abstract

**Background:** Long-term persistent symbiotic associations may result in co-speciation and can be inferred if species trees of hosts and symbionts are congruent in topology and divergence times. Co-speciation has been seen to occur relatively frequently in obligate associations, but is less common in parasitic or facultative ones, mainly due to the difference in horizontal transmission rates. The long-term vertical inheritance and close host association of obligate endosymbionts also generally result in smaller genomes than in facultative endosymbionts. Here, we investigate co-speciation and genome reduction using highly similar strains of the endosymbiont *Wolbachia* infecting *Drosophila* species from the willistoni and saltans groups, where only one strain, *w*Pau, infecting *D. paulistorum*, is obligate.

**Results:** We sequenced the *Wolbachia* genomes from five species of the willistoni and saltans groups and constructed phylogenies. Topological congruence was found between these *Wolbachia* strains and the nuclear DNA of their hosts, except for *w*Pau and *D. paulistorum*, but full topological congruence was observed between *Wolbachia* and the host mitochondrial DNA. However, assuming temporal congruence, we estimated extremely low evolutionary rates in *Wolbachia* of 10^-10^-10^-11^ changes/site/year. Additionally, the obligate *w*Pau strain was found to have a larger genome than closely related facultative strains, mainly due to an ongoing expansion of an IS4 element. Furthermore, *w*Pau has lost a large proportion of its prophage WO genes, but the *cif* genes, known to be involved in the CI phenotype, are intact. Finally, nine of the eleven genes from the prophage WO-associated Undecim cluster are uniquely duplicated.

**Conclusions:** The congruent topologies between *Wolbachia* and their willistoni and saltans group hosts indicate co-speciation. However, the high similarity between *Wolbachia* strains, which results in low mutation rate estimates, challenges this interpretation. Contrary to the expectations of the genome reduction theory, we observed an increase in genome size in the obligate *w*Pau strain, potentially driven by a decreased population size. Finally, the duplication of the Undecim cluster despite a major loss of other prophage-associated genes suggests that the genes in the Undecim cluster are under strong selection and potentially play a role in the obligate association between wPau and their *D. paulistorum* hosts.

## Background

Bacterial symbionts are ubiquitous among animals but vary widely in phenotypic effects, level of dependency, and evolutionary persistence within host lineages. Mutualistic symbioses often involve tight associations with the host that may become obligate and thereby evolutionary persistent in a host lineage over time. In contrast, parasitic or pathogenic symbionts are usually facultative and evolutionarily short-lived unless they can manipulate their host to ensure their transmission across generations, by so-called reproductive parasitism (Zeng and Wiens 2021). Hence, the evolutionary persistence of a symbiont typically suggests that it is either an obligate mutualist or a reproductive parasite. The phenotypic effect, dependency, and persistence of a symbiotic association are thus interconnected and have been shown to influence how symbiont genomes evolve (McCutcheon and Moran 2012; Fisher et al. 2017). Numerous studies (Wernegreen 2015; McCutcheon and Moran 2012; Bobay and Ochman 2017) have demonstrated that obligate symbionts typically have smaller genomes, fewer pseudogenes, repeats, and mobile elements than facultative symbionts. The reduced genome size in obligate symbionts reflects their long-term reliance on the host, with the loss of non-essential gene functions and ongoing deletions facilitated by genetic drift. In contrast, facultative symbionts, which maintain a more transient association with their hosts, retain larger genomes that provide greater metabolic versatility and independence as well as more repeats and mobile elements.

*Wolbachia* is an intracellular symbiont that infects many different arthropod species as well as some nematodes (Kaur et al., 2021) and forms both facultative and obligate host associations. It is mainly vertically transmitted via the maternal line to the next generation (Russell et al. 2019) but is also frequently horizontally transmitted between host species (Vancaester and Blaxter 2023). *Wolbachia* can affect many aspects of its host’s biology, including fecundity, immunity, and behaviour (Kaur et al. 2021). However, it is best known in arthropods for its ability to alter host reproduction. One such reproductive phenotype is cytoplasmic incompatibility (CI), where infected males are unable to produce offspring with uninfected females. The reproductive advantage thereby gained by infected females over uninfected females, can result in the spread of *Wolbachia* into a host population (Shropshire et al. 2020) as well as persistence in a population once the infection has reached a high frequency (Turelli and Hoffmann 1995). Even so, only a handful of examples of evolutionary persistent *Wolbachia* are currently known. Phylogenetic congruence, which is a hallmark of evolutionary persistence and co-speciation, has, for example, been observed between obligate mutualistic *Wolbachia* in filarial nematodes (Lefoulon et al. 2016) and bed bugs (Balvín et al. 2018), but also in non-obligate associations between *Wolbachia* and *Nasonia* wasps where *Wolbachia* induces CI (Raychoudhury et al. 2008) and *Nomada* solitary bees (Gerth and Bleidorn 2016) where the phenotypic effect of *Wolbachia* is unknown. However, phylogenetic congruence between host mitochondrial DNA and *Wolbachia* is regularly observed for closely related host species, suggesting that the cytoplasmic co-inheritance of mitochondria and *Wolbachia* is frequently maintained at least over shorter times (Charlat et al. 2009; Ilinsky 2013; Zhang et al. 2013; Cariou et al. 2017; Bakovic et al. 2020).

*Wolbachia* genomes generally conform to the genome reduction theory. Facultative *Wolbachia* strains infecting various arthropods have genomes ranging between 1.1-1.8 Mb in size (Werren et al. 2008; Vancaester and Blaxter 2023) with a substantial fraction of mobile elements and other repeats and relatively many pseudogenes. Obligate mutualistic *Wolbachia* of nematodes have smaller genomes, ranging from 0.5-1.0 Mb, with few pseudogenes and a low fraction of mobile elements or repeats (Lefoulon et al. 2020; Dudzic et al. 2022). Additionally, prophage WO regions are present in nearly all *Wolbachia* genomes except in *Wolbachia* of nematodes, where they are absent or highly degraded (Gavotte et al.

2007; Bordenstein and Bordenstein 2022). This is important since prophage WO contains genes responsible for reproductive parasitism, such as *cifA* and *cifB* responsible for CI (Beckmann et al. 2017; LePage et al. 2017) and *wmk* involved in male-killing (Perlmutter et al. 2019), as well as the Octomom region affecting *Wolbachia* titer and virulence (Chrostek and Teixeira 2015; Duarte et al. 2021). Furthermore, the Eukaryotic Association Module (EAM) of prophage WO encodes several proteins suggested to play a role in *Wolbachia*-host interactions (Bordenstein and Bordenstein 2016).

In the Neotropical *Drosophila* of the willistoni and saltans groups, most tested species carry *Wolbachia* with high similarity to each other and the well-known *w*Au strain from *Drosophila simulans* (hereafter called *w*Au*-*like) (Miller and Riegler 2006). The high frequency of infections and similarity between the *Wolbachia* strains found in the two *Drosophila* groups indicates that their ancestor may have been infected with a *w*Au*-*like *Wolbachia* and that the two partners may have been co-speciating. However, this observation is puzzling, as co-speciation suggests evolutionary persistence and almost all the *Wolbachia* strains in willistoni and saltans group flies are believed to be facultative (Müller et al. 2012) and, in some cases, unable to induce CI and/or lack the *cif* genes (Martinez et al. 2015, Martinez et al. 2021). Only the *Wolbachia* strain *w*Pau infecting *D. paulistorum* from the willistoni group, a superspecies consisting of multiple reproductively isolated semispecies, is considered obligate mutualistic, as previous results have shown that all collected *D. paulistorum* flies are infected. Additionally, complete removal of *w*Pau by antibiotic treatment causes lethality and reduces the *w*Pau titer, resulting in distorted ovary development (Miller et al. 2010), thus affecting fecundity. Additionally, *w*Pau is likely involved in the ongoing host speciation of *D. paulistorum* as it contributes to assortative mating between different semispecies of *D. paulistorum* (Miller et al. 2010), affects pheromone profiles (Schneider et al. 2019) and contributes to the distinct gene expression pattern in heads and abdomens of three semispecies (Baião et al. 2019). *D. paulistorum* is also known to carry two distinct mitotypes, α and β, and multiple mitochondrial introgressions with related Neotropical *Drosophila* species likely occurred during divergence of its semispecies, as indicated by discordant mitochondrial and nuclear phylogenies (Baião et al. 2023). Thus, if a potential mitochondrial donor species carried *Wolbachia*, *w*Pau could have entered *D. paulistorum* via introgression.

The Neotropical *Drosophila* system thus offers the opportunity to investigate evolutionary persistence and co-speciation in non-obligate *Wolbachia* associations and study genomic differences between very closely related facultative and obligate *Wolbachia* strains. To do so, we sequenced and assembled *Wolbachia* genomes from five *Drosophila* species: *D. paulistorum, D. willistoni* and *D. tropicalis* from the willistoni group, and *D. prosaltans* and *D. septentriosaltans* from the saltans group and inferred whole genome phylogenies. We found congruence between four of five *Wolbachia* strains and host nuclear genes and full congruence between *Wolbachia* and host mitochondrial genomes.

Additionally, we discovered that the genome of the obligate *Wolbachia* strain *w*Pau is larger than closely related facultative strains due to the expansion of IS4 elements. However, the *w*Pau genome also had features commonly associated with ongoing genome reduction, like a lower coding percentage and more pseudogenes. Finally, despite a reduced prophage WO content, we found the WO-associated *cif* genes intact and a unique duplication of the Undecim cluster, suggesting they are under selection and thus potentially important for the symbiosis between *w*Pau and its host.

## Materials and Methods

### DNA Extraction and Sequencing

*Drosophila* lines used in this study and the sequencing data produced from each of them are summarized in Tables S1 and S2. For *D. paulistorum* O11, total DNA for Oxford Nanopore (ONT) sequencing was extracted from many adult flies of both sexes using the MagAttract HMW DNA Kit (Qiagen) to obtain high molecular weight DNA. For Illumina sequencing, total DNA was obtained as described in Baiaõ et al 2023. For *Wolbachia* strains *w*WilP98, *w*ProPE2, *w*PauFG111, *w*PauPOA1 and *w*PauTP37, DNA samples used for PacBio sequencing were enriched for *Wolbachia* using a differential centrifugation and filtration procedure to obtain a *Wolbachia* cell pellet, which was then subjected to whole genome amplification using the Repli-g midi kit (Qiagen) as described in (Ellegaard et al., 2013). For *w*WilP98 and *w*ProPE2, Illumina sequences enriched for *Wolbachia* were obtained using the same protocol.

The ONT library was produced and sequenced using an R9.4 PromethION flow cell at the Uppsala Genome Center, Uppsala, Sweden. Basecalling was performed with Guppy 4.3.4 and the HAC model. Illumina TruSeq libraries were produced and sequenced at the Uppsala SNP and Seq platform on an Illumina 2500 HiSeq machine generating 2x125 bp reads. The Illumina reads were quality filtered and trimmed using Trimmomatic-0.30 with parameters –phred33 ILLUMINACLIP:illumina_adapter.fasta:2:40:15 LEADING:3 TRAILING:3 SLIDINGWINDOW:4:15 MINLEN:95 (Bolger et al., 2014) and error-corrected using SPAdes-3.5.0 --sc --only-errorcorrection (Bankevich et al., 2012). For PacBio, 5 kb SMRTbell libraries were created. Each library was run with P6-C4 chemistry in one SMRT cell on an RSII PacBio instrument at Uppsala Genome Center.

### Genome Assemblies

*Wolbachia* genomes were assembled using several types of data and pipelines. For *w*PauO11, reads from total DNA sequencing (i.e. including the *Drosophila* host) were filtered using Filt-long v0.2.1 with quality priority and the parameters “--target_bases 12500000000 - -min_length 1000 --mean_q_weight 10” and assembled with Nextdenovo v2.5.0 using correction options: read_cutoff = 1k, genome_size = 250m, sort_options = -m 7g -t 2, minimap2_options_raw = -t 3, pa_correction = 17, correction_options = -p 2 and assemble options: minimap2_options_cns = -t 3, nextgraph_options = -a 1. The produced assembly contained only one *Wolbachia* contig that was subsequently polished with Illumina reads Pilon v1.24 (Walker et al. 2014) with the --fix indels parameter and circularized. The genomes of the two strains *w*WilP98 and *w*ProPE2 were assembled using Illumina and PacBio reads from amplified DNA enriched for *Wolbachia*. The Illumina reads were assembled using Spades v3.15.5 (Bankevich et al. 2012) with kmer sizes 97,103,107,113 and the PacBio reads were used for scaffolding with the --pacbio parameter. Illumina and PacBio reads were mapped against the assemblies using BWA-mem (Li and Durbin 2009) with default parameters for Illumina reads and using PacBio settings for PacBio reads to check the correctness of the genome assembly. Overlapping contigs were merged in Consed (Gordon et al. 1998). See Table 1 for assembly statistics of complete genomes. For all other assemblies, BWA-mem v0.7.17-r1198-dirty with default parameters was used to map Illumina reads on the *w*PauO11, *w*ProPE and *w*WilP98 genomes and a set of concatenated *Wolbachia* genomes representing the *Wolbachia* diversity (81 genomes from supergroups A, B, C, D, E, F, L and M, Table S3). Mapped reads were filtered using samtools v.1.13 (Danecek et al. 2021) view with the -f 1 -F 12 parameters to include only pairs where both were mapped. Filtered reads were then subsampled to approximately 100X with seqtk v1.3-r117-dirty sample -s 100 before assembly. The reads were then assembled using Spades v3.15.5 with kmer sizes 97,103,107,113, and the *w*PauFG111, *w*PauPOA1 and *w*PauTP37 assemblies were scaffolded with PacBio reads using the flag -pacbio.

**Table 1.** Genomic features in complete *w*Au-like and outgroup genomes.

|  | wPauO11<sup>1</sup> | wWilP98<sup>1</sup> | wProPE2<sup>1</sup> | wAu<sup>2</sup> | wInn<sup>3</sup> | wMel<sup>4</sup> | wCin2<sup>5</sup> |
| --- | --- | --- | --- | --- | --- | --- | --- |
| **Genome size(bp)** | 1,365,198 | 1,268,527 | 1,339,466 | 1,268,455 | 1,290,587 | 1,267,782 | 1,538,351 |
| **GC (%)** | 34.8 | 35.2 | 35.2 | 35.2 | 35.3 | 35.2 | 35.1 |
| **CDSs** | 1000 | 1016 | 1033 | 996 | 1073 | 1011 | 1473 |
| **% Coding** | 73.8 | 77.5 | 76.3 | 75.7 | 70.1 | 76.0 | 86.2 |
| **Pseudogenes** | 177 | 123 | 130 | 122 | 227 | 111 | 216 |
| **Repeats (%)** | 22.9 | 11.4 | 15.8 | 11.0 | 16.8 | 9.8 | 13.6 |
| **Phage WO (%)** | 8.8 | 10.8 | 12.6 | 10.6 | 9.0 | 9.5 | 24.3 |
| **IS elements (%)** | 15.3 | 7.5 | 9.4 | 7.6 | 9.9 | 7.0 | 7.6 |

<sup>1</sup>*Wolbachia* genomes from Neotropical *Drosophila* sequenced in this study, <sup>2</sup>Baião GC et al., 2021, <sup>3</sup>Hill T et al., 2022, <sup>4</sup>Wu M et al., 2004, <sup>5</sup>Wolfe TM et al., 2021.

The Illumina assemblies were then filtered by removing contigs that deviated from the mean sample distribution in kmer coverage and GC content. The kmer coverage of the contigs was determined from the fasta headers in the output from Spades. In cases where kmer coverage had a bimodal distribution, the minimum between the two peaks was identified using the quantmod package in R (Ryan and Ulrich 2022) and used as threshold. Contigs with kmer coverage below this threshold were blasted against the assembled contigs and were removed if the sequence existed. This procedure ensured that redundant low coverage contigs were removed. Remaining low coverage contigs were blasted against additional *Wolbachia* genomes and proteins. Contigs with a blast alignment length longer than 100 aa and percent identity over 60%, or blast alignment length longer than 200 nucleotides and percent identity over 70%, were kept. Any contig with an extreme GC value (outside the 99% interval of a normal distribution of GC) was removed. Finally, all remaining contigs were searched with Kraken against a custom database (containing a selection of archaea, bacteria, plasmids, viruses, human, *D. melanogaster* and *D. willistoni* sequences) and the contigs with a match to *Drosophila* were removed. All assemblies were polished once using Pilon with the Illumina reads used for assembly.

### Genome Annotation

For the complete genomes, *w*PauO11, *w*ProPE2 and *w*WilP98, Prokka v1.14.6 (Seemann 2014) with the parameters --genus Wolbachia --compliant --rfam --addgenes --usegenus was used for annotation. Next, Pseudofinder v1.1.0 (Syberg-Olsen et al. 2022) with parameters -ce and -db with a database containing all *Wolbachia* proteins from NCBI (2022- 03-02), was used to mark potential pseudogenes. Additionally, hmmsearch as implemented in pfam_scan.pl was used for domain prediction with the PFAM database (Bateman et al.2002). The Illumina reads used for the assemblies were mapped back to each genome with BWA-mem to verify frameshifts and nonsense mutations. All marked pseudogenes were inspected manually and annotated as pseudogenes if the start or stop codons were missing, they contained nonsense or frameshift mutations or were truncated by, for example, an IS element insertions.

The draft assemblies were annotated the same way, except that 1) Pseudofinder was run with length_pseudo set at 0.8 and the protein database only included *Wolbachia* genomes *w*Au, *w*Ha, *w*Ma, *w*No, *w*San, *w*Tei, *w*Yak, *w*PauO11, *w*ProPE2 and *w*WilP98 that were manually annotated either in this or previous studies (Ellegaard et al. 2013; Baiaõ et al. 2021), 2) only pseudogene calls in CDS sequences were inspected manually, 3) pseudogenes that we identified in the complete genomes (*w*PauO11, *w*ProPE2 and *w*WilP98), but were not found by pseudofinder in the draft genomes, were transferred to the draft genome annotation if they matched with high similarity and extensive coverage.

For identifying repeated sequences, nucmer from the MUMmer 4.0 package (Marçais et al. 2018) was used with flags --maxmatch --nosimplify to align each genome to itself. The positions of non-overlapping repeat bases were then extracted using a perl script. ISEScan v 1.7.2.3 (Xie and Tang 2017) was used to identify IS elements. Bar plots of the percentage of repeats and IS elements in each genome were made in R (R Core Team 2018; Wickham et al. 2019). Phage WO regions were annotated by blastp searches to *Wolbachia* genomes (*w*Mel, *w*Au, *w*Ha, *w*Ma, *w*No, *w*San, *w*Tei, *w*Yak) with previously identified phage WO regions. All genomic regions that contained similarities to known phage WO genes were then manually inspected, and borders were determined based on Bordenstein and Bordenstein 2022.

### MLST phylogeny of *Wolbachia* genomes

Genomes that contained the keyword “Wolbachia” in the taxon name, were downloaded from NCBI using the command line tool “datasets” (https://github.com/ncbi/datasets) on May the 6^th^ 2022. tblastn of the Blast v2.7.1+ suite (Camacho et al. 2009) with the parameters -- max_hsps 1 --evalue 1e-10 was used to search for the five *w*Mel MLST proteins and Wolbachia surface protein (Wsp) (GenBank accession numbers AAS13898.1, AAS14036.1, AAS14201.1, AAS14415.1, AAS14719.1, AAS14884.1) in 1504 downloaded genomes.

Genomes with blast hits for all six genes were kept for phylogenetic inference. Nucleotide sequences of the identified genes were extracted from each genome using samtools faidx, and translated, aligned with MAFFT v6.850b using the linsi algorithm (Katoh and Toh 2008), pruned to remove gap sites present in more than 50% of aligned sequences and then backtranslated to nucleotides again. A phylogeny of the six genes from 1262 genomes was then constructed using RaxML v8.2.9 (Stamatakis 2014) with parameters –f a –m GTRGAMMA –x 12345 –p 12345 -# 100. The resulting phylogenetic tree as visualized in Figure 1 and all other figures presenting phylogenetic trees (Fig. 2, Fig. 4B, Fig. S1, Fig. S2, Fig S3, Fig. S5 and Fig. S6) were made using FigTree v1.4.4 (Rambaut 2009).

**Figure 1.**
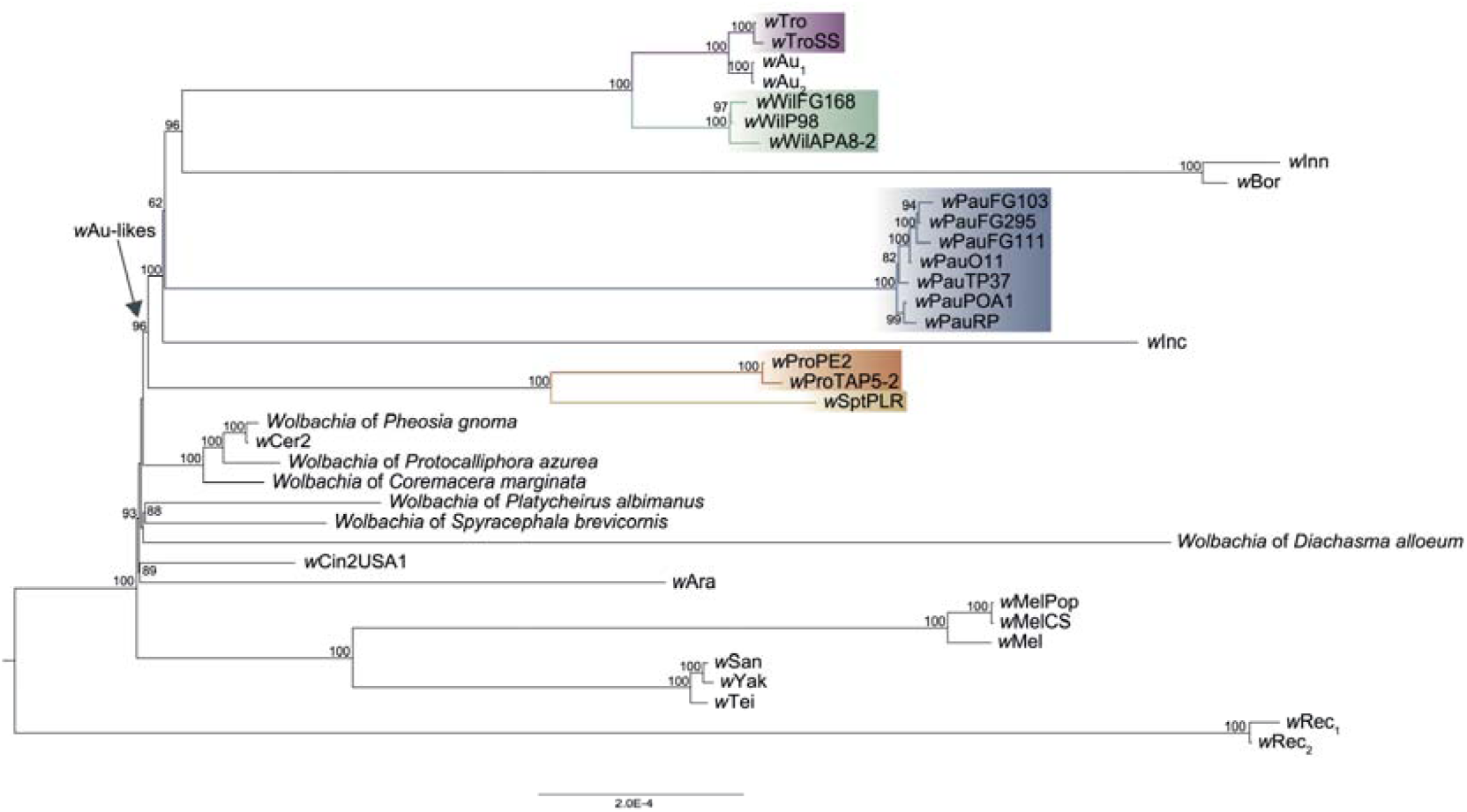
Phylogeny of *Wolbachia* genomes with high similarity to *w*Au-like_ws_ strains. The branches with *Wolbachia* strains associated with *Drosophila* species from the willistoni and saltans groups, i.e., the *w*Au-like_ws_ *Wolbachia* strains, are coloured. The clade referred to as the *w*Au-like is indicated with an arrow. The tree was inferred from an alignment of 620 single-copy orthologs with IQ- TREE. Support values from 1000 ultrafast bootstrap replicates are displayed on the nodes of the tree. Only bootstrap values greater than 60 are shown.

**Figure 2.**
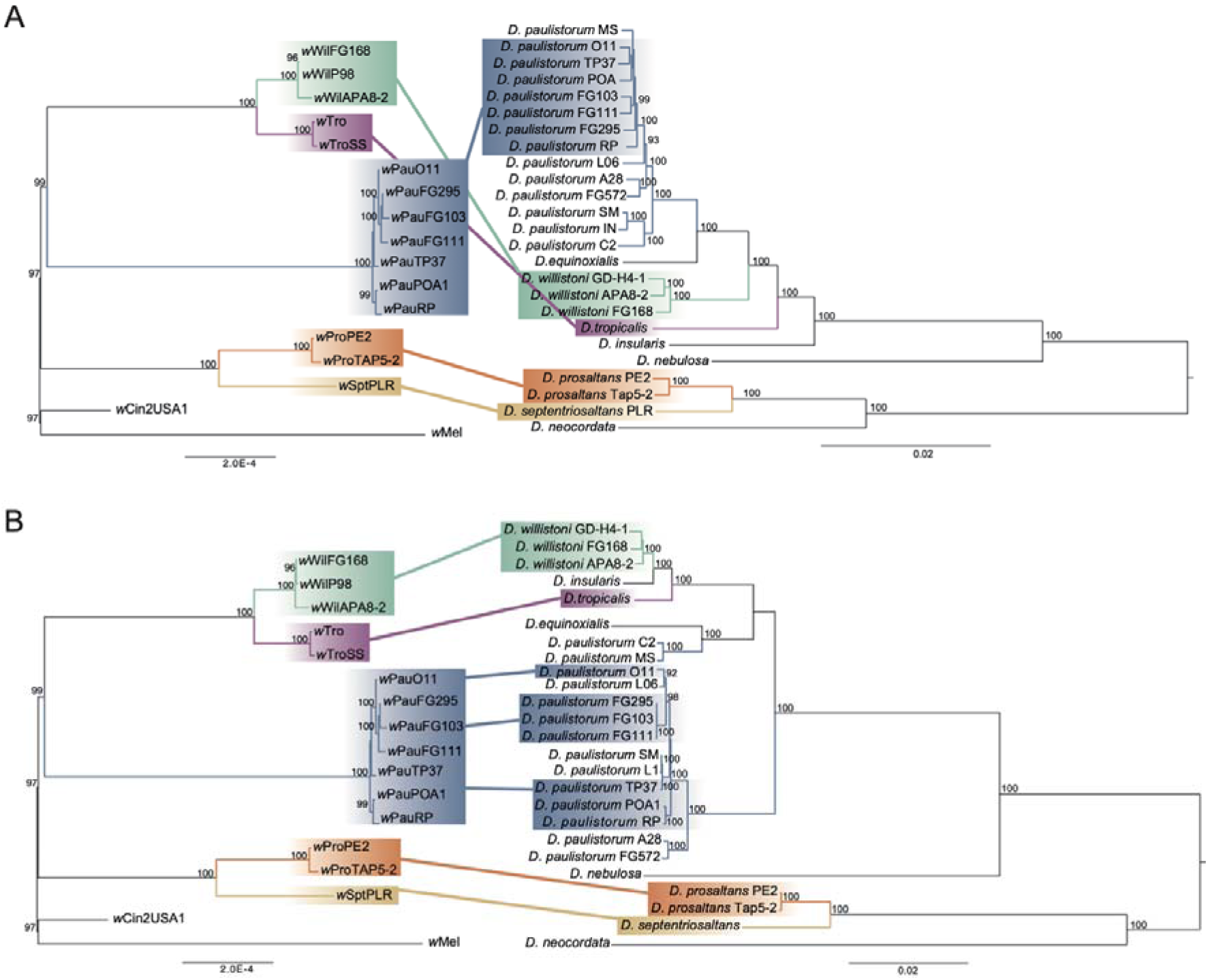
Phylogenetic comparison between *w*Au-like *Wolbachia* and *Drosophila* of the willistoni and saltans groups. A) Maximum Likelihood Phylogenies of *w*Au-like *Wolbachia* on the left and 692 nuclear-encoded proteins of the host on the right (from Baião et al. 2023). The *w*Au-like *Wolbachia* phylogeny is a subset of the tree in Figure 1. B) Maximum Likelihood Phylogenies of *w*Au- like *Wolbachia* on the left (same as in A) and the whole mitochondrial genome of the neotropical *Drosophila* host species on the right (from Baiaõ et al. 2023).

**Figure 3.**
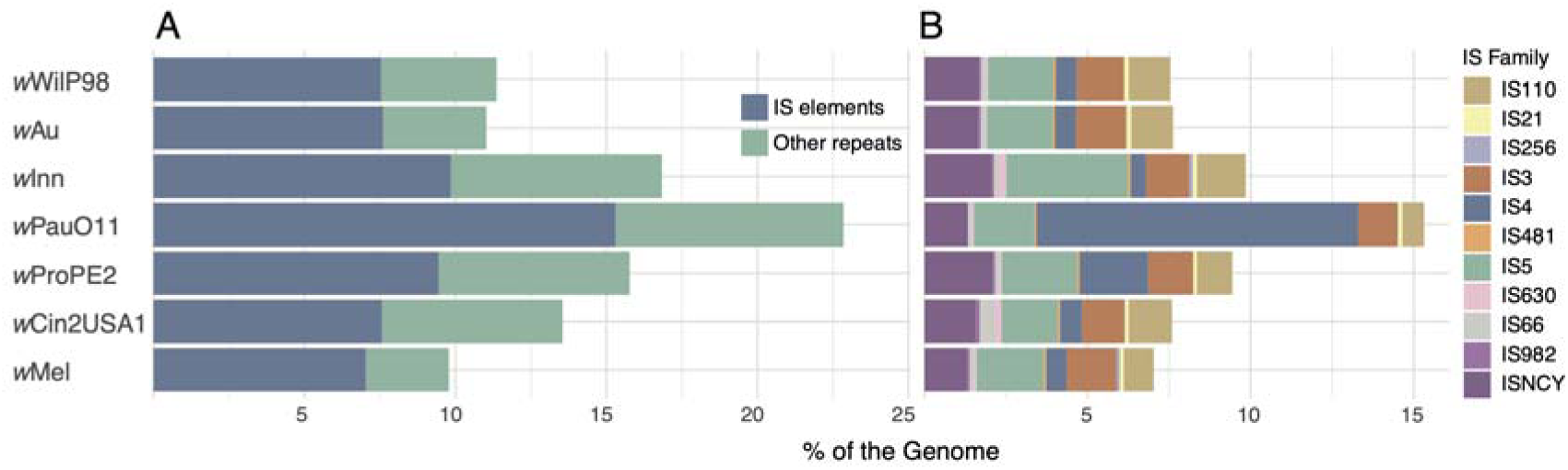
Repeats and IS-elements in complete genomes of *w*Au-like strains. A) Percentage of repeats in each genome. B) Percentage of each IS family in the genomes, as identified by ISEscan.

**Figure 4.**
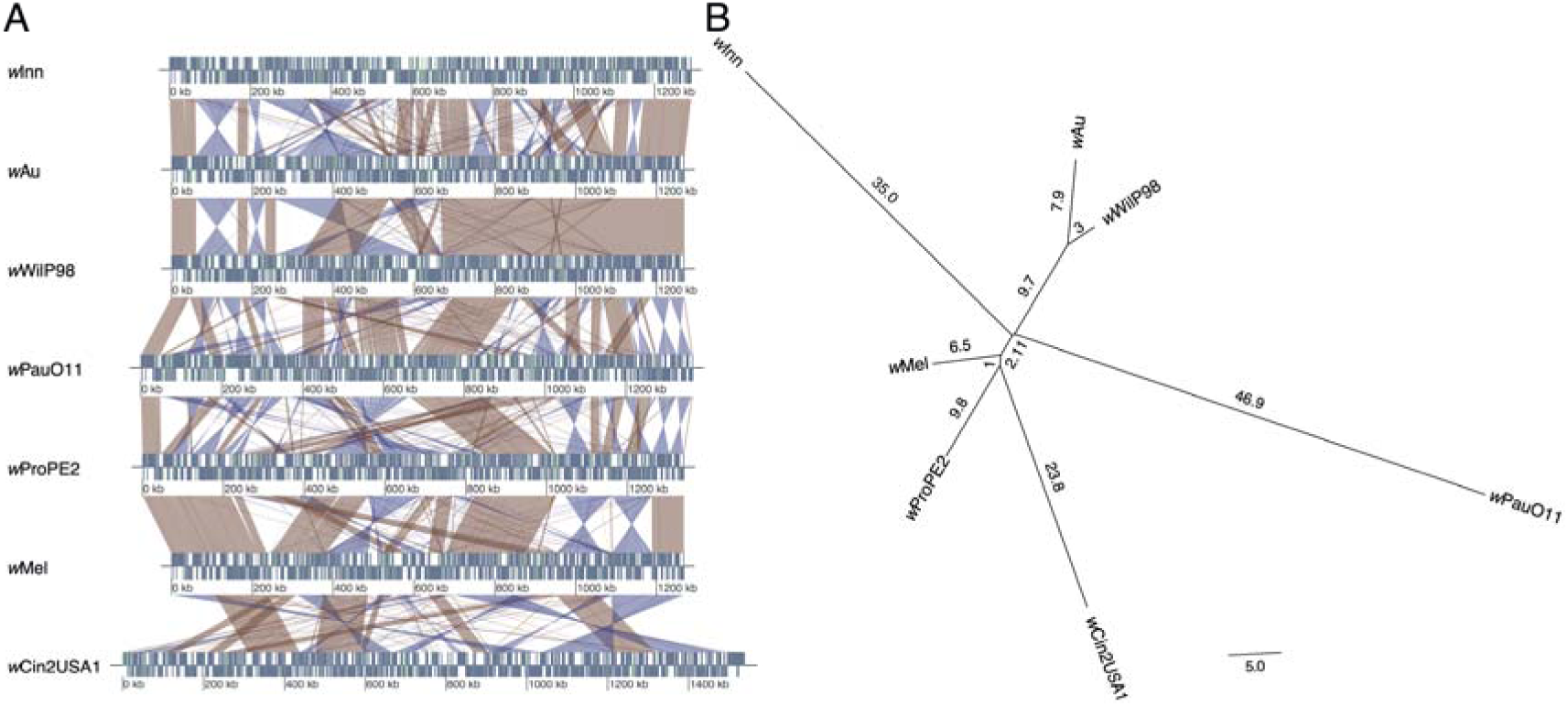
Genomic rearrangements in *w*Au-like strains. A) Whole genome alignments of complete *Wolbachia w*Au-like genomes and outgroup. For each genome, genes are visualized with blue and pseudogenes with green rectangles on either the forward or reverse strand (above and below the midline respectively). Similarities between genomes are visualized as red and blue ribbons, with red representing forward and blue reverse hits. B) Gene order phylogeny of the same genomes as in A. Branch lengths written on each branch represent the estimated number of genomic rearrangements.

### Protein clustering and strain phylogeny

At the time of downloading *Wolbachia* genomes, the ones from Vancaester and Blaxter 2023 were not yet available. Hence, to include them in our analysis, we downloaded the genomes classified as supergroup A by the authors. We then annotated the genomes from all strains classified as supergroup A *Wolbachia* in the MLST phylogeny and by Vancaester and Blaxter 2023 as described above for our draft assemblies, used their proteomes for clustering and inferred a phylogeny from single-copy genes (Table S4). Clustering was done with Orthofinder v2.5.4 (Emms and Kelly 2019) with -I 3 and filtered all-against-all blastp results as input. Blastp hits between two matching proteins were filtered so that the size of one should not be smaller than 60% of the size of the other, the e-value should be lower than 10^-5^ and the alignment length should be at least 80% of the size of the smaller protein. The nucleotide sequences of genes from orthogroups with single-copy proteins were extracted, translated, aligned with mafft-linsi, pruned to remove gap sites present in more than 50% of the aligned sequences and backtranslated to obtain codon-based nucleotide alignments.

To ensure that the genes used in the phylogeny did not show signs of recombination, we ran Phipack (Bruen et al. 2006) with the -p 100000, -o, -w 50 on all alignments. Phipack runs three tests to detect recombination, the Phi test, the max χ2 test and the NSS test. For a gene to be considered as recombined, at least two of the three recombination tests needed to have a p-value lower than 0.05. Single-copy gene alignments without recombination were then concatenated with geneStitcher.py from Utensils (https://github.com/ballesterus/Utensils/blob/master/geneStitcher.py).

IQ-TREE v2.2.0 (Minh et al. 2020) was then used for phylogenetic inference of the species tree using the concatenated alignments of the single-copy genes, a partition per gene, and the parameters -m MFP, -B 1000 -T 20 --seed 12345. Individual gene trees were inferred using the same parameters. The phylogenetic trees were visualized using FigTree v1.4.4. To test if the data fit better with the placement of *w*Pau according to the host nuclear phylogeny rather than our strain phylogeny, we followed Shen et al. 2017 and used IQ-TREE v1.6.12 with the same parameters as in the strain phylogeny together with the -z flag to add the two topologies (shown in Figure S3) in Newick format and the -wsl and -wpl flags to get the likelihood for each site and gene. The differences in the gene likelihoods between the two topologies were calculated and plotted in R (R Core Team 2018; Wickham et al. 2019). To measure the gene (gCFs) and site (sCFs) concordance factors, we used the same pruned concatenated phylogeny as in the log-likelihood differences and we produced pruned gene trees including the same strains with preserved branch lengths using a Python script and the ete3 library (Huerta-Cepas et al. 2016). To calculate the gene and site concordance factors, the pruned species tree was then provided to IQ-TREE with the flag -t, the fasta alignments with -s, the partition file with -p, the gene trees were provided with --gcf and --scf 100 was used.

### Substitution rates

Pairwise synonymous substitution rates were calculated with codeml from the PAML v4.8 package (Yang 2007) using alignments of the 620 single-copy *Wolbachia* genes from the ortholog clustering (Table 2), as well as the alignments of 692 BUSCO genes and 13 mitochondrial genes from *Drosophila* (the same as in Baião et al 2023) generated the same way as the *Wolbachia* alignments (see above). For substitution rate estimates from MCMCTree, we calibrated our phylogeny based on estimates from Suvorov et al. 2022 on the willistoni-saltans group divergence at around 14.2-20.6 million years and the *D. willistoni*-1. *D. tropicalis* divergence at around 4.2-6.1 million years. A rough estimate of 19-21 million years for the age of the tree and a time unit of 100 million years gives a beta of 156 for the gamma distribution of rates. We used alpha parameters of 1, 5, and 10 for the shape of the distribution to describe scenarios of low, medium, and high mutation rates. The whole alignment of single-copy genes as one partition was given as input (ndata=0, seqtype==0), and the GTR substitution model (model=8) was used with independent rates (clock=2). Each of the three rate scenarios was run two times to verify convergence of estimates for the chains, for 20,100,000 generations (burnin=100,000, sampfreq=1,000, nsample=20,000).

**Table 2.**
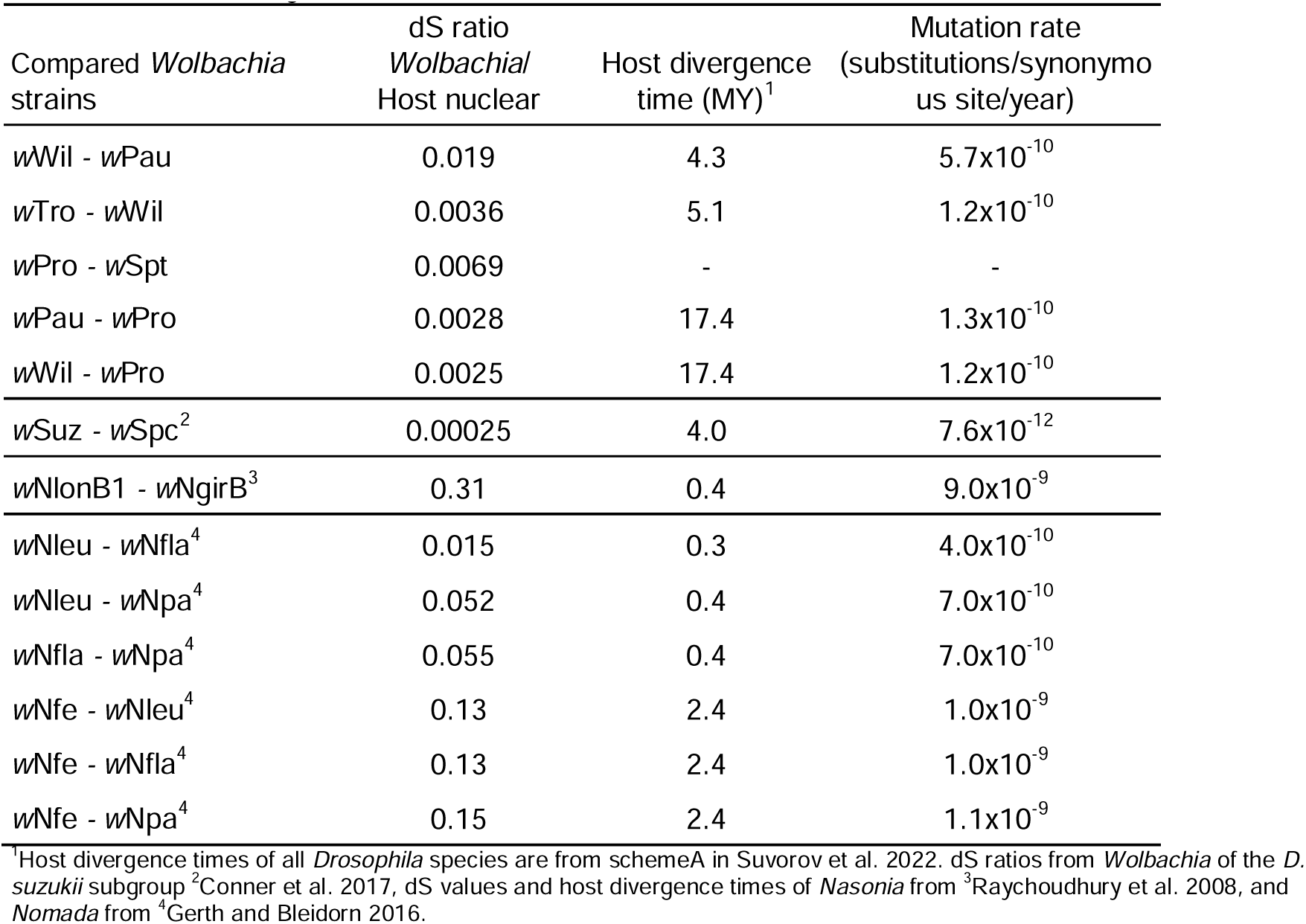
Evolutionary rates of *Wolbachia*.

| Compared *Wolbachia* strains | dS ratio<br>*Wolbachia*/<br>Host nuclear | Host divergence<br>time (MY)<sup>1</sup> | Mutation rate<br>(substitutions/synonymous site/year) |
| --- | --- | --- | --- |
| wWil - wPau | 0.019 | 4.3 | 5.7x10<sup>-10</sup> |
| wTro - wWil | 0.0036 | 5.1 | 1.2x10<sup>-10</sup> |
| wPro - wSpt | 0.0069 | - | - |
| wPau - wPro | 0.0028 | 17.4 | 1.3x10<sup>-10</sup> |
| wWil - wPro | 0.0025 | 17.4 | 1.2x10<sup>-10</sup> |
| wSuz - wSpc<sup>2</sup> | 0.00025 | 4.0 | 7.6x10<sup>-12</sup> |
| wNlonB1 - wNgirB<sup>3</sup> | 0.31 | 0.4 | 9.0x10<sup>-9</sup> |
| wNleu - wNfla<sup>4</sup> | 0.015 | 0.3 | 4.0x10<sup>-10</sup> |
| wNleu - wNpa<sup>4</sup> | 0.052 | 0.4 | 7.0x10<sup>-10</sup> |
| wNfla - wNpa<sup>4</sup> | 0.055 | 0.4 | 7.0x10<sup>-10</sup> |
| wNfe - wNleu<sup>4</sup> | 0.13 | 2.4 | 1.0x10<sup>-9</sup> |
| wNfe - wNfla<sup>4</sup> | 0.13 | 2.4 | 1.0x10<sup>-9</sup> |
| wNfe - wNpa<sup>4</sup> | 0.15 | 2.4 | 1.1x10<sup>-9</sup> |

<sup>1</sup>Host divergence times of all *Drosophila* species are from schemeA in Suvorov et al. 2022. dS ratios from *Wolbachia* of the *D. suzukii* subgroup <sup>2</sup>Conner et al. 2017, dS values and host divergence times of *Nasonia* from <sup>3</sup>Raychoudhury et al. 2008, and *Nomada* from <sup>4</sup>Gerth and Bleidorn 2016.

### Gene order phylogeny

Syntenic blocks between the complete genomes listed in Table 1 were obtained using SynChro_linux (January 2015) (Drillon et al. 2014) with a delta value of 2. The output from SynChro was then used for phylogenetic reconstruction with PhyChro (downloaded July 2023 from http://www.lcqb.upmc.fr/phychro2/) (Drillon et al. 2020).

### IS4 element and Undecim cluster copy numbers in draft *w*Pau genomes

To estimate the number of IS4 elements and Undecim cluster copies in the draft *w*Pau genomes, we first masked all copies of the IS4 element and one copy of the Undecim cluster in the *w*PauO11 genome. To do that, we created an IS4 element consensus sequence by blasting (blastn) one IS4 element from *w*PauO11 against the *w*PauO11 genome, extracting the sequences for all matches, aligning them with MAFFT v7.490 with flags “--maxiterate 1000” and “--localpair” and building a consensus with Jalview v2.11.4.0 (Waterhouse et al. 2009). The consensus IS4 element sequence was then blasted against the *w*PauO11 genome using blastn and all hits were masked. The Undecim cluster copies were first identified by homology to the Undecim cluster genes in the *w*Mel genome. One of the Undecim cluster copies, at positions 898.960-913.092 in *w*PauO11, was then manually masked. The Illumina reads used to assemble the draft *w*Pau genomes were then mapped to the masked *w*PauO11 genome and the IS4 element consensus sequence using BWA-mem v0.7.17-r1198-dirty. Repeated areas were identified from the annotation (see above) and non-repeated regions of the *w*PauO11 genome were identified using “bedtools complement” and the repeated regions as input. Coverage over the IS4 element consensus sequence and the unmasked Undecim cluster copy was calculated with “samtools coverage” and the coverage of the non-repeated regions of the genome was calculated by getting the average per-base coverage in these regions by using “samtools depth”. The copy numbers of the IS4 element and the Undecim cluster were calculated by dividing the average coverage over these regions by the average coverage over the non-repeated regions of the genome.

### SNP phylogeny of *w*Pau genomes

To get a more resolved relationship between the *w*Pau strain variants, we called SNPs for all *w*Pau strain variants from the Illumina reads mapped to the masked *w*PauO11 genome (see above). Reads with MQ<1 were filtered out and bcftools was used to call SNPs using the command “bcftools mpileup --threads 6 -Ou -d 1000 -f $genome -b bam.file.list.txt | bcftools call --threads 6 -cv --ploidy 1 -Ob -o variants_cflag.bcf”. To filter low-quality variants, the command “bcftools view --types snps -i ‘QUAL>30 & INFO/DP>10 & MQ>30’” was used.

The variants located in non-repeated areas of the *w*PauO11 genome were extracted using “bcftools view” with the coordinates identified as described above and “bcftools consensus” was used to create pseudo-chromosomes for each strain. The sequence for each pseudo- chromosome was put into a multi-fasta file and used as input for IQ-TREE v2.2.0 using the same parameters as in the *Wolbachia* strain phylogeny. The phylogeny was visualized using FigTree v1.4.4.

### IS4 element insertion dynamics

To detect differences in IS4 insertions between the *w*Pau genomes, we identified improperly mapped read pairs in the Illumina reads that were mapped to the masked *w*PauO11 genome (see above). If an IS4 element is present in the genomes with mapped reads, the insert size of paired reads in this region is 0, as one read of the pair would map on the masked *Wolbachia* genome and the other read would map on the consensus IS4 sequence. To get such improperly paired reads, we filtered with samtools view using parameters -F2, MQ>0 and -e “tlen==0”. Reads that mapped within 750 bases upstream of the start or 750 bases downstream of the end of an IS4 element in wPauO11 (as specified in a provided bed file) are from an IS4 element shared with *w*PauO11. Reads mapping at other places are associated with IS4 insertions not present in *w*PauO11, and they were filtered by using samtools view with the -U -L flag and the same bed file as above. If an IS4 element is present in *w*PauO11 but absent in the genome with mapped reads, the reads from a pair are further apart than expected, specifically the insert size would appear to be larger than the ca. 1500 bases of the IS4 element, and both reads of the pair should map on the masked *Wolbachia* genome. To get such improperly paired reads, reads mapping 750 bases upstream and 750 bases downstream of IS4 elements (as above) and that had an insert size larger than 1500 bases but shorter than 5000 bases were filtered using samtools view -e “(tlen>1500 && tlen<5000) || (tlen>-5000 && tlen < -1500)”. The filtered read alignments of improper pairs were visualized in IGV (v2.16.0) to identify unique IS4 element insertions across the different *w*Pau strain variants. The *w*Pau whole genome SNP phylogeny was then used to infer where and when IS4 copies were inserted in the *w*Pau strain variants.

Because complete deletions of IS4 elements occur rarely, differences were explained by insertions, rather than deletions. Additionally, we inferred the number of IS4 element copies present at each node of the tree using complete *Wolbachia* genomes. We identified the shared IS4 elements across the *w*Au-like strains, *w*Mel and *w*Cin2USA1 by blasting (blastn v2.7.1+) genomes to each other and visualizing the regions flanking IS4 elements, identified by ISEscan, with Artemis comparison tool (ACT) (Carver et al. 2005). An IS4 element was considered absent from a genome if a region had the same gene order as an IS4 element-containing region of another genome but didn’t contain an IS4 element or if an IS4 element-containing region in other genomes shared no synteny with the target genome.

### Gene content variation between genomes *w*Au-like *Wolbachia*

To compare gene content across the different *Wolbachia* genomes, we used the gene count output of Orthofinder to produce a hierarchical clustering of the genomes based on gene presence/absence. First, we excluded all single-copy genes and then we assigned the count of multi-copy genes to 1 so that we cluster based on presence and not copy-number. Then, we used the “agnes” function for agglomerative clustering with the “average” method from the cluster v2.1.4 package in R (Maechler et al. 2021).

Additionally, the variability in gene content between the *w*Au-like_ws_ genomes was investigated by identifying orthogroups where only one or several of the lineages, *w*Pau, *w*Tro/*w*Wil and *w*Pro/*w*Spt, were present and one or several lineages were absent (Table 3, Table S5 and Table S6). Unless otherwise stated, presence was counted if a gene was present in the complete genome of the lineage (*w*PauO11, *w*WilP98 and *w*ProPE2) and at least one draft genome of the same lineage.

**Table 3.**
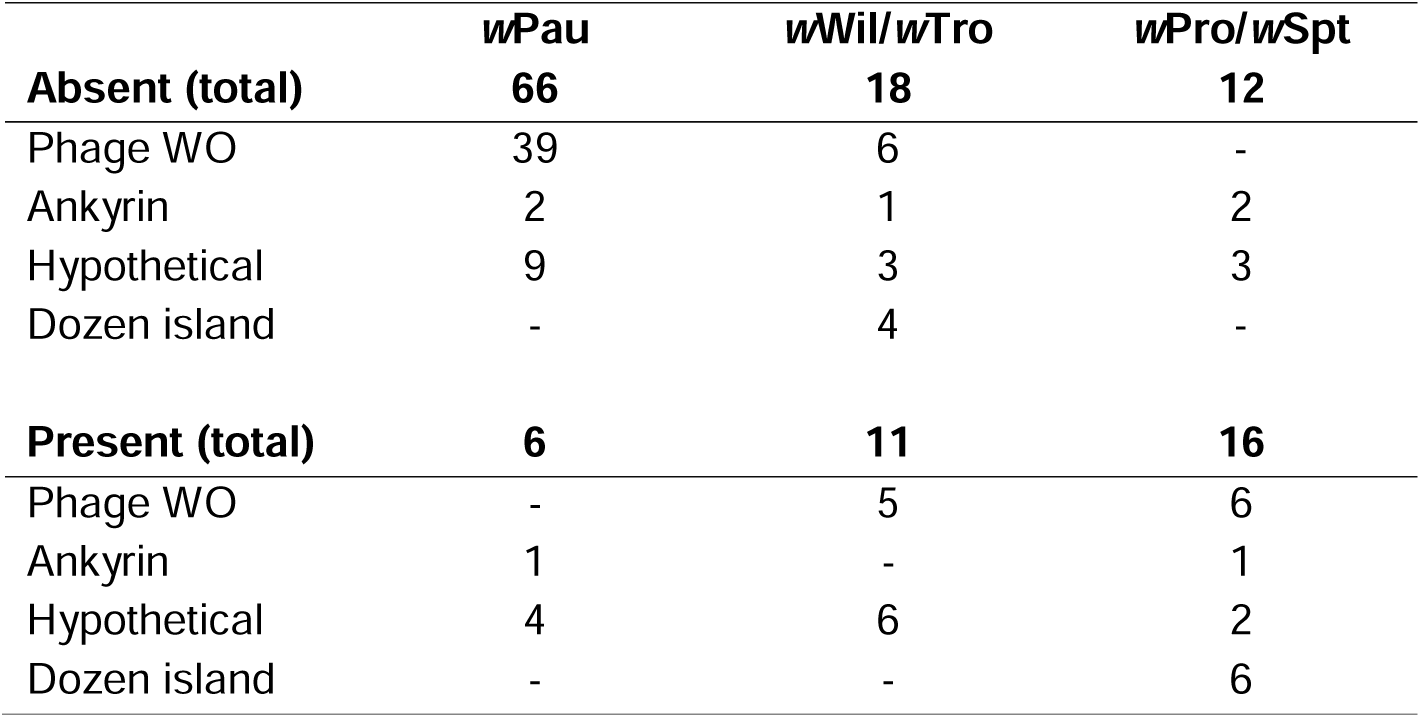
Gene content variation between *w*Au-like_ws_ *Wolbachia*. The numbers represent the presence or absence of an orthogroup in one lineage compared to the other two lineages in the table.

All figures presenting genome and gene comparisons (Fig. 4A, Fig. 7, Fig. S7 and Fig. S8) were made with genoPlotR (Guy et al. 2010), after using blastn to search each genome against every other genome.

### Prophage WO classification and gene module assignment

To investigate the prophage WO content in detail, we identified prophage WO regions in the complete *w*Au-like genomes, *w*PauO11, *w*ProPE2, *w*WilP98, *w*Inn, *w*Au, and in *w*Mel, *w*Cin2USA1. All proteins of prophage WO origin were re-clustered to orthogroups with OrthoFinder v2.5.4 with -I=1 and custom blastp (as above), which put most phage WO paralogues into single orthogroups. We manually validated the orthogroups based on the functional annotation and gene order in complete genomes (Table S7). The orthogroups containing proteins with the same functional annotation that were syntenic were manually merged. For orthogroup, we inferred a phylogeny using IQ-TREE v2.2.0 as described for single-copy genes. Genes were assigned to WOa and WOb based on whether they formed a clade with either WOMelA or WOMelB genes and/or were syntenic with WOMelA or WOMelB. For orthogroups where a WOMelB gene was present, we followed the classification of Bordenstein and Bordenstein 2022. Otherwise, we looked at what module the neighboring genes belonged to and assigned the orthogroup to that module. If an orthogroup had no functional annotation and could not be assigned to a module using the above method, it was assigned to the EAM module if flanked by genes in a structural module on one side and an EAM gene on the other side.

## Results

We assembled three complete (*w*PauO11, *w*ProPE2 and *w*WilP98) and eleven draft genomes (Table 1, Table S1) from *Wolbachia* strains infecting three different species of the *Drosophila* willistoni group: seven *w*Pau strain variants from the Orinocan (OR) semispecies of *D. paulistorum* (as determined by the host phylogeny), three *w*Wil strain variants from *D. willistoni,* one *w*Tro strain variant from *D. tropicalis,* and two species of the *Drosophila* saltans group: two *w*Pro strains variants from *D. prosaltans* and one *w*Spt strain variant from *D. septentriosaltans* (Table S2). We refer to each strain variant by the *Wolbachia* strain name followed by the *Drosophila* line it was sequenced from, i.e., *w*PauO11 is the *w*Pau strain variant from the *D. paulistorum* line O11.

### Co-speciation between *Wolbachia* and their Neotropical *Drosophila* hosts

To determine if *Wolbachia* co-speciated with their willistoni and saltans group hosts, we started by producing a phylogeny of our new genomes and other closely related *Wolbachia* genomes. To achieve that, we downloaded all publicly available *Wolbachia* genomes and inferred a phylogeny using the five Multi-Locus Strain-Typing (MLST) genes (Baldo et al. 2006) and the *wsp* gene. We determined 23 of the 1613 downloaded genomes to be closely related to ours (Table S4) and clustered their proteomes with the proteomes of our 14 genomes. The resulting 620 clusters with single-copy orthologous genes, all without detectable recombination, were used to produce a phylogeny.

The concatenated gene alignment contained only 0.55% parsimoniously informative sites (3,088 of 562,755 sites), showing the very close relationship between all 37 strains.

Despite this, the inferred phylogeny had high support for a clade including the five *Wolbachia* strains from the willistoni and saltans *Drosophila* groups*, w*Tro, *w*Wil, *w*Pau, *w*Pro and *w*Spt (Figure 1). This clade, which also contained *Wolbachia* strain *w*Au from the old-world host *D. simulans*, further included *Wolbachia* strains *w*Inn, *w*Bor and *w*Inc, which infect *Drosophila* species of various other new-world species groups. Thus, from here on, these nine highly similar *Wolbachia* strains are what we refer to as *w*Au-like, and the strains infecting willistoni and saltans group flies are referred to as *w*Au-like_ws_. Finally, as eight of the nine *Drosophila* hosts are new-world species, we assume that *w*Au-like infections evolved in Americas and have just recently infected old-world hosts like *D. simulans*.

The concatenated gene alignment of these nine *w*Au-like genomes contained only 0.3% informative sites and 13.2% of the genes had no informative sites at all. Even so, inside the *w*Au-like clade, all strain and most strain variant relationships, except the position of *w*Pau (bootstrap 62), were highly supported.

Thus, to evaluate the position of *w*Pau, we calculated the gene (gCF) and site concordance factors (sCF) for the internal branches of the subtree that only included one representative genome (*w*PauO11, *w*WilP98, *w*Tro, *w*ProPE2 and *w*SptPLR) of each *w*Au- like_ws_ strain, which are the focus of the co-speciation analysis. The concordance factors represent the percentage of gene trees, gCF, or informative sites, sCF, that support a split in the tree over all other possible splits. After removing single-copy genes without any informative sites, we found that almost all informative sites (99.6-100%) place *w*PauO11 between the *w*Wil/*w*Tro and *w*Pro/*w*Spt clades and most of the genes (76.1-81.2%) also agree with this placement (Figure S1). Thus, the position of *w*PauO11 in our inferred phylogeny had strong support from informative sites and a majority of the individual genes. Furthermore, the low bootstrap value for the split between *w*Pau and other strains only occurred when the *w*Inc strain of *D. incompta* was included in our phylogenetic analysis.

When *w*Inc was excluded, the *w*Pau strain variants were in the same position in the tree but with high bootstrap support (100) (Figure S2). Since our focus is on the topology of the *w*Au- like_ws_ *Wolbachia* strains, we also inferred phylogenies using only those genomes and an outgroup. The resulting phylogeny confirms that the position of *w*Pau is supported (bootstrap support 99) with respect to the other *w*Au-like_ws_ *Wolbachia* strains (Figure 2, left part).

When comparing our inferred *w*Au-like_ws_ strain phylogeny to the nuclear phylogeny of the willistoni and saltans group hosts (Baiaõ et al. 2023), we saw that the trees were congruent except for the position of *w*Pau and *D. paulistorum* (Figure 2A). Rather than being positioned inside the clade of the other two willistoni group strains, *w*Wil and *w*Tro, *w*Pau is 19 positioned between the willistoni and saltans group (*w*Pro and *w*Spt) strains. However, full congruence was observed when comparing the *Wolbachia* tree with the phylogeny of mitochondrial genomes from the same *Drosophila* species (Figure 2B) (Baiaõ et al. 2023).

To test if our *Wolbachia* data fit better with the host nuclear or mitochondrial topologies, we calculated the log-likelihood difference per gene between topologies congruent with either the mitochondrial or the nuclear phylogeny. We found that most *Wolbachia* genes strongly supported the mitochondrial-like topology, and a few genes weakly supported the nuclear-like topology (Figure S3). Overall, our results indicate that the *w*Au-like_ws_ *Wolbachia* strains are highly congruent with the host mitochondrial phylogeny, and partly congruent with the nuclear phylogeny of the hosts.

For co-speciation to be true, the divergence times of the hosts and *Wolbachia* strains should also match (referred to as temporal congruence). To get an idea about temporal congruence, we calculated relative pairwise synonymous substitution rates of *w*Au-like_ws_ *Wolbachia* and host genes (Table S8) and compared them to previously described cases of *Wolbachia* co-speciation in *Nasonia* and *Nomada* (Raychoudhury et al. 2008, Gerth and Bleidorn 2016) and horizontal transmission in the *D. suzukii* subgroup (Conner et al. 2017). The relative substitution rates ranged from 0.00025 for *Wolbachia* in the *D. suzukii* subgroup to 0.31 for *Wolbachia* of *Nasonia* (Table 2), with *w*Au-like_ws_ *Wolbachia* in between (0.0025 - 0.019).

We also calculated mutation rates for these *Wolbachia* strains by dividing the pairwise synonymous substitution rates by the divergence times of their hosts (if available). The estimated mutation rates for *w*Au-like_ws_ *Wolbachia* were between 1x10^-10^ to 5x10^-10^ synonymous substitutions per synonymous site per year, which is in the same range as some of the *Wolbachia* in *Nomada,* but again higher than in *Wolbachia* from the *D. suzukii* subgroup, where *Wolbachia* was suggested to be horizontally transmitted and lower than the ones in *Nasonia,* where *Wolbachia* is suggested to co-speciate with the hosts (Table 2, Table S8). Hence, the evolutionary rates of *w*Au-like_ws_ did not match those from previously suggested examples of *Wolbachia* co-speciation or horizontal transmission.

Additionally, we estimated the mutation rate of *w*Au-like_ws_ *Wolbachia* in a Bayesian framework using host divergence times for the willistoni-saltans groups and *D. willistoni* – *D. tropicalis* (Suvorov et al. 2022) as priors. This resulted in a mutation rate for *w*Au-like_ws_ *Wolbachia* between 2.6 - 6.8x10^-11^ substitutions per site per year (Table S9), which is one order of magnitude smaller than the rate based on synonymous substitutions.

### Repeat expansion and rearrangements in the genome of obligate strain *w*Pau

Contrary to the expectations for an obligate strain, the *w*PauO11 genome was larger than all other *w*Au-like *Wolbachia* genomes (Table 1), had a lower coding content and 25-50% more repeats than both the genomes of *w*Au-like *Wolbachia* and two closely related facultative outgroup strains, *w*Mel and *w*Cin2USA1 (Table 1, Figure 3A). Specifically, we discovered a unique and extensive expansion of an IS4 element, which accounted for 9.9% of the total genome size of *w*PauO11 (Figure 3B).

Since repeats act as templates for genomic rearrangements via intragenomic homologous recombination, we examined gene order differences between our closely related genomes. From a whole genome alignment, we saw that the *w*PauO11 genome exhibited many changes in gene order when compared to other *w*Au-like genomes (Figure 4A). To quantify this, we identified pairwise syntenic blocks between all genomes and used this information to construct a phylogeny (Figure 4B). In the resulting tree, *w*PauO11 was on the longest branch, indicating that it has experienced more genomic rearrangements than the other *Wolbachia* genomes.

These findings highlight that the *w*PauO11 genome differs significantly from the genomes of other *w*Au-like_ws_ strains in several ways, such as having a high proportion of non-coding DNA and repeats and frequent genomic rearrangements. Interestingly, a similar pattern, except for the large repeat expansion, is also seen in the genome of the *w*Au-like strain *w*Inn from *D. innubila*.

### IS4 expansion is still ongoing in the *w*Pau genomes

Given the significant expansion of IS4 elements in *w*PauO11, we investigated whether these elements are still actively transposing in *w*Pau genomes. By identifying all complete and incomplete copies in the *w*PauO11 genome, we estimated that it originally had at least 96 complete IS4 elements (Figure S4). To quantify IS4 elements in all other *w*Pau genomes, we masked all but one copy in the *w*PauO11 genome, mapped the Illumina reads of each *w*Pau strain to it and calculated the coverage over the IS4 element relative to non-repetitive regions. Applying this approach to Illumina reads from *w*PauO11, we estimated that this genome harboured 98 IS4 copies, which closely matched our observation of 96 copies.

When extended to other *w*Pau variants, the analysis revealed variability in copy numbers of the IS4 element across genomes (95-124 copies/genome) (Table S10). Additionally, by examining improper read pairs, we identified the positions of IS4 element insertions that occurred after the divergence of the *w*Pau strain variants (Table S11). To get a better resolution of the *w*Pau strain variant relationships and thus be able to calculate the number of insertions on each branch more correctly, we inferred a phylogeny based on whole genome SNPs between the *w*Pau variants (Figure S5). Using this tree topology, we identified seven IS4 element insertion events that occurred after the divergence of the *w*Pau strain variants (Table S11, Figure S6). Hence, both read mapping approaches indicated that the IS4 element is still actively transposing in these *w*Pau genomes.

Next, we approximated the rate and timing of IS4 element insertions in all complete genomes of the *w*Au-like clade. Using copy number counts, synteny comparisons and our phylogeny, we estimated that the genome of the ancestor of all *w*Au-like strains had four IS4 elements (Table S12) and the number of IS4 element insertions in all non-*w*Pau genomes to be 22 (Figure S6). Using branch lengths as an approximation of time, we calculated a rate of 6,571 insertions/(substitution/site) for the non*-w*Pau genomes. We inferred 88 IS4 element insertions on the branch leading up to the ancestor of the *w*Pau variants by estimating a minimum of 93 IS4 element copies in the genome of the last common ancestor of *w*Pau.

This gave us a rate of 123,422 insertions/(substitution/site). The seven IS4 element insertions in the *w*Pau strain variants gave us an estimated rate of 83,333 insertions/(substitution/site). The insertion rate of IS4 elements is thus 18.8 times greater in the branch leading to the *w*Pau ancestor and 12.7 times greater in the current *w*Pau than in the other analysed *Wolbachia* strains (Table S13). Our analyses thus showed that the IS4 element insertion rate was slightly higher in the *w*Pau genome in the past than it is today, but expansion is still ongoing with a relatively fast rate.

### Major loss of prophage WO genes, but retention of CI genes and duplication of the Undecim cluster in *w*Pau

To explore which genes might be linked to phenotypic differences among *w*Au-like_ws_ strains, we clustered the genomes based on the presence and absence of genes.

Our analysis revealed that the genomes of *w*Pro and *w*Spt clustered together, as did those of *w*Wil, *w*Tro, and *w*Au, consistent with their phylogenetic relationships (Figure 1). However, the gene content of *w*Pau was notably distinct from other the *w*Au-likews genomes (Figure 5).

**Figure 5.**
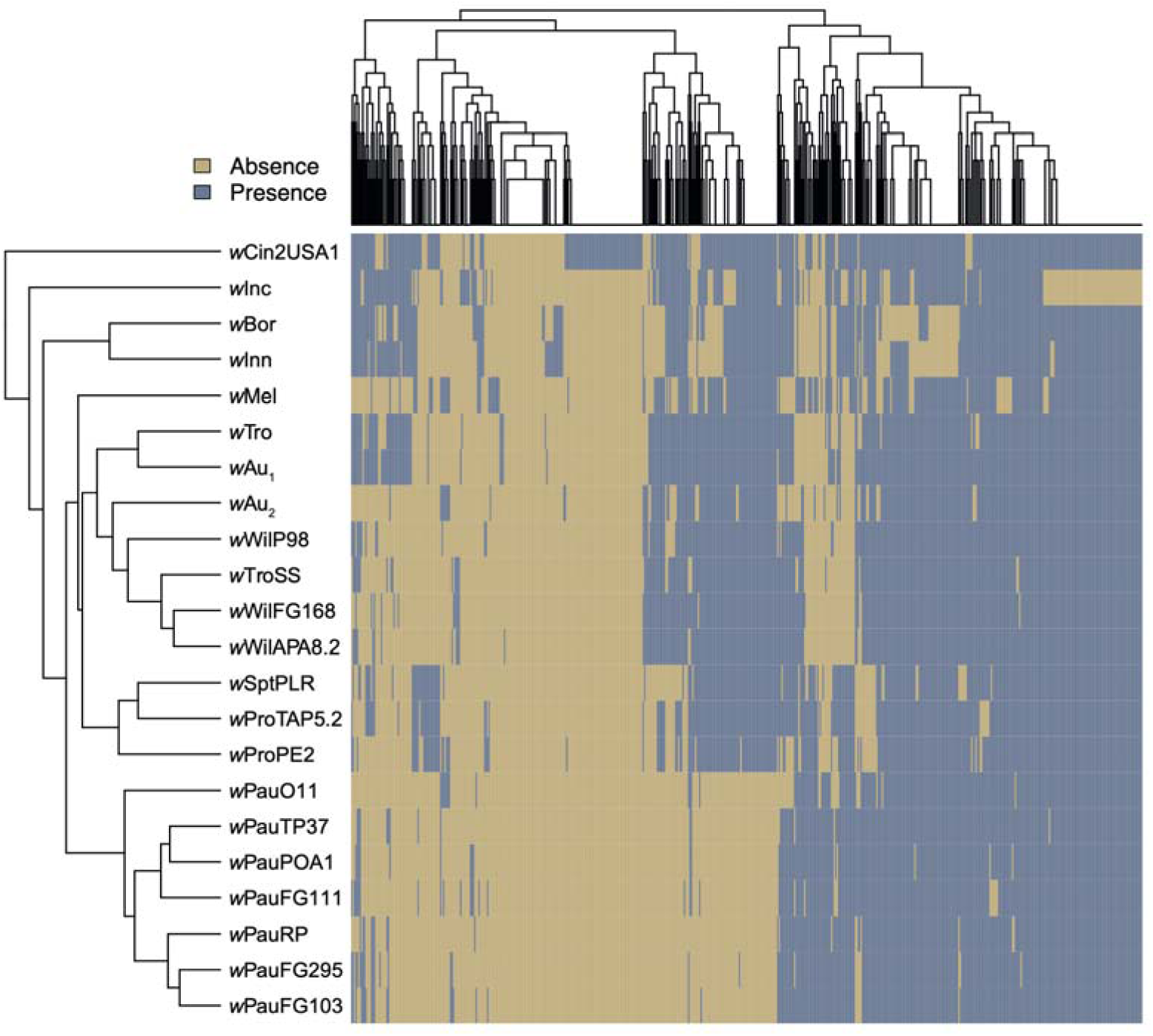
Gene content clustering of *Wolbachia w*Au-like genomes. The cladogram on the left shows how the *Wolbachia* strains cluster based on the presence and absence of orthogroups in each genome, which is depicted in the plot to the right.

We analysed the variation in gene content between the *w*Au-like_ws_ strains further, by identifying which genes were uniquely present or absent in the three lineages, *w*Pau, *w*Wil/*w*Tro and *w*Pro/*w*Spt. Our results showed that *w*Pau (defined as *w*PauO11 and at least one *w*Pau draft genome) lacked many genes compared to the other *w*Au-like_ws_ strains (Figure 5, Table S5), rather than having unique ones (Table 3, Table S6). Most missing genes were from prophage WO regions (Table 3), as suggested by the lower proportion of prophage WO sequences in the *w*PauO11 genome. Of the 39 missing prophage WO- associated genes, most encoded structural phage WO proteins, such as tail, head, or baseplate proteins, whereas only six belonged to the Eukaryotic Association Module (EAM).

Given the large number of missing prophage WO genes in *w*PauO11, we wanted to determine if they all came from the same prophage WO copy. The prophage WO regions identified in the complete *w*Au-like genomes were highly similar to either the WO-A or WO-B regions in *w*Mel (Wu et al. 2004) and are referred to here as WOa and WOb, respectively.

By classifying the prophage WO proteins into WOa- and WOb-like, we found that WOa-like prophages are generally more degraded than WOb-like prophages in all genomes, with the Lysis, Tail, and Tail fiber modules of the former entirely missing in all *w*Au-like genomes as well as in *w*Mel and *w*Cin2USA1 (Figure 6).

**Figure 6.**
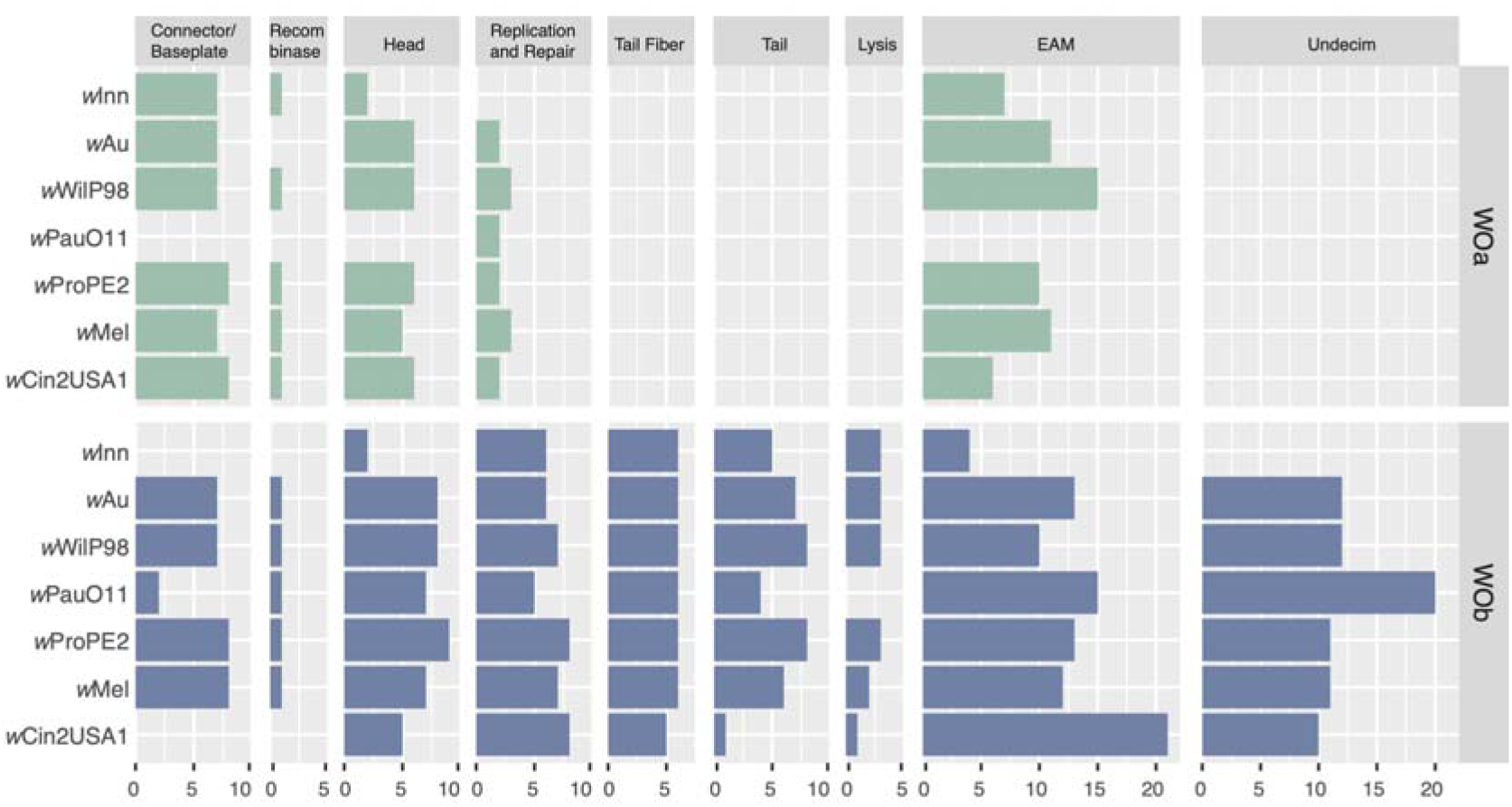
Prophage WO genes in complete *w*Au-like genomes and outgroup. Number of genes per module for the two prophage WO variants identified in *w*Au-like and related genomes. WOa genes were found in the same clade as genes from WO-A in *w*Mel and WOb genes were found in the same clade as genes from WO-B in *w*Mel when inferring single gene phylogenies.

**Figure 7.**
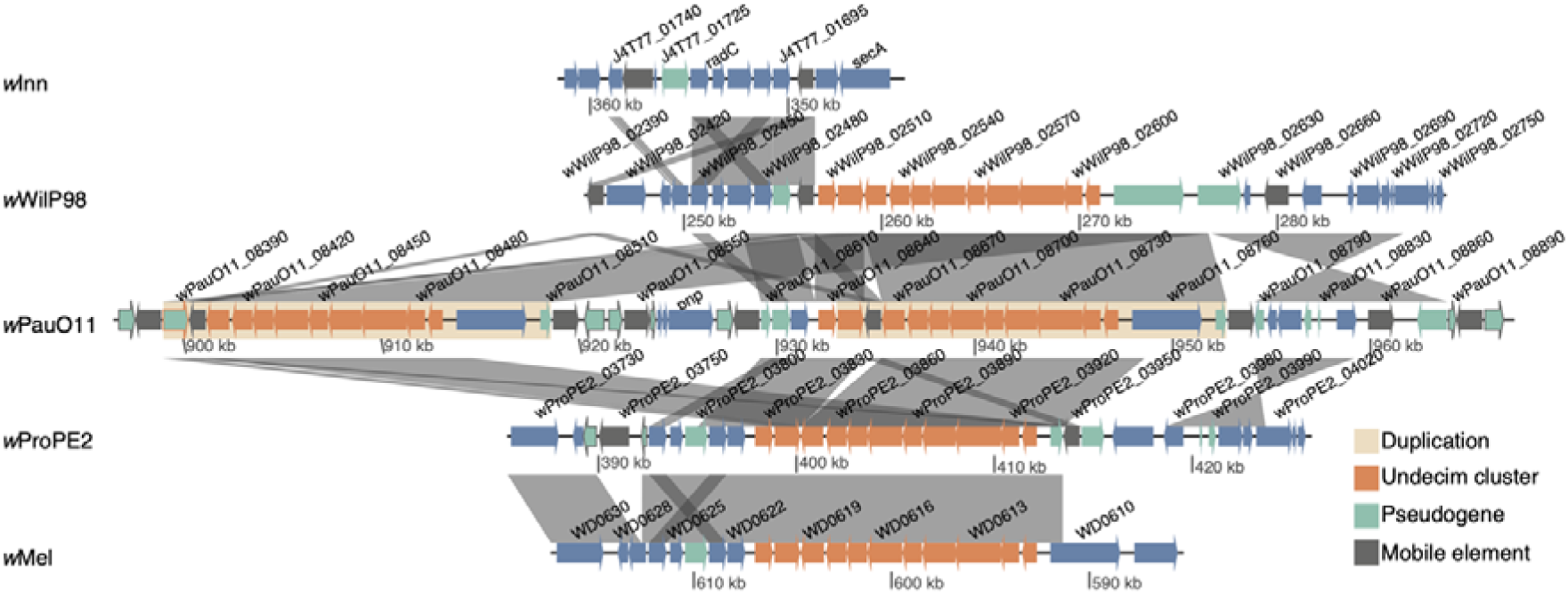
Duplication of the phage WO-associated genes in the Undecim cluster in *w*PauO11. Similarities between genomes are visualized with grey bands and the duplication is marked in yellow. Genes in orange are part of the Undecim cluster. Mobile elements are dark grey and pseudogenes green, with the border colour indicating Undecimcluster or mobile element.

However, in *w*PauO11, the WOa-like prophage was almost completely absent, except for a few genes within the Replication and Repair module. The WOb-like prophage in *w*Pau also displayed a reduced gene content, particularly in the Connector/Baseplate module. Conversely, the *w*Inn genome, which also has a low proportion of prophage WO, retained all Baseplate/Connector genes in WOa but instead lost them in WOb. The presence of genes associated with both WOa and WOb in *w*Pau and other complete *w*Au-like genomes suggests that intact copies were likely present in their common ancestor. Thus, *w*Pau has predominantly lost genes from the WOa copy, but both prophage WO copies have experienced gene loss.

Apart from prophage WO genes, nine ankyrin repeat domain-containing genes (ANKs) and nine hypothetical proteins were absent from all *w*Pau strain variants but present in the *w*Wil*/w*Tro and *w*Pro*/w*Spt lineages. Of the remaining 11 protein clusters absent in all *w*Pau, functions varied and included two non-WO phage proteins, a Fic protein, Type IV secretion system protein VirB6, Malonyl-CoA decarboxylase, Cytochrome d ubiquinol oxidase, subunit I and Exopolysaccharide synthesis ExoD-related protein. All missing protein clusters in *w*Pau were found in *w*Au-like genomes from one or several of the non-*w*Au-like_ws_ strains, indicating that they have likely been lost in *w*Pau rather than gained in the other *w*Au-like_ws_ lineages.

Much fewer protein clusters were uniquely absent in the other two *w*Au-like_ws_ lineages. Twelve clusters were missing from the *w*Pro and *w*Spt genomes, and 20 were absent in *w*Wil and *w*Tro (Table 3). Noteworthy, the *w*Wil/*w*Tro lineage lacks the prophage WO-associated genes responsible for CI, *cifA* and *cifB*, as previously observed by Martinez et al. 2021. Additionally, we found that *cifB* contains a frameshift mutation in the *w*Spt genome, likely rendering it non-functional (Figure S7, Table S14). Furthermore, the *w*Bor and *w*Inn strains also have frameshift mutations in *cifB* as well as non-sense mutations in *cifA*, while *w*Inc has a non-sense mutation in *cifB* but an intact *cifA*. Thus, the only *w*Au-like strains that potentially can induce CI are *w*Pro and *w*Pau. Since highly similar Type I *cif* genes are found in all *w*Au-like genomes except in one clade (*w*Tro/*w*Wil/*w*Au), the common ancestor of *w*Au-likes likely carried Type I *cif* genes. Hence, our results indicate that the *cif* genes have been lost or pseudogenized independently multiple times in the *w*Au-like clade.

Only six protein-coding genes, encoding four hypothetical proteins, one ANK and a ComF family protein, were present in *w*PauO11 (and at least one other draft *w*Pau genome) but absent in the other *w*Au-like_ws_ strains (Table 3). Potentially, one of the hypothetical proteins and the ANK gene were acquired by *w*Pau, since no homologs were found in any other *w*Au-like genome. Additionally, one of the hypothetical proteins is homologous to WD0462 in *w*Mel, which correlated with CI induction in another set of *Wolbachia* strains (Baião et al. 2021). However, since the gene is pseudogenized in *w*Pro, a strain known to induce CI, the correlation was not seen among our strains. Finally, the ComF family protein, with a potential role in DNA uptake, is pseudogenized in other *w*Au-like_ws_ genomes.

More genes are uniquely present in the *w*Wil/*w*Tro and the *w*Pro/*w*Spt lineages. For example, homologs of WD0463, the neighbouring gene of WD0462, are only intact in *w*Pro/*w*Spt. This genomic region showed additional variability, with *w*Pau and *w*Pro carrying an extra gene between WD0462 and WD0463 (Figure S8). Even though the function of either of these genes is unknown, the region is highly variable between *Wolbachia* genomes (Baião et al. 2021), even among these very close relatives. Additionally, we noted that six genes in *w*Pro and five in *w*Spt from the Dozen Island, a genomic region likely stemming from an integrated plasmid (Baião et al. 2021, Martinez et al. 2022), were not present in *w*Pau or *w*Wil/*w*Tro. An additional four genes of the Dozen Island were found in *w*Pro/*w*Spt and *w*Pau but lost from the wWil/wTro lineage.

Finally, although the loss of prophage WO genes appeared to be frequent in the *w*Pau genome, we identified a large and unique duplication of a prophage WO-associated region in *w*PauO11 containing ten of the eleven genes in the Undecim cluster (Bordenstein and Bordenstein 2022) (Figure 7).

To determine if the duplication was present in the other *w*Pau strain variants, we estimated the copy number by mapping reads from those genomes to the *w*PauO11 genome with one copy masked. Coverage over the Undecim cluster was twice as large as coverage of non-repeated sequences for *w*PauFG111, *w*PauPOA1 and *w*PauTP37 (Table S10), indicating duplication of Undecim in those genomes. For *w*PauFG103, *w*PauFG295 and *w*PauRP, coverage over the Undecim cluster was more like in non-repeated sequences. However, for those lines, the standard deviation of coverage for non-repeats was as much as 30% of the mean, compared to 10% for the other lines. This suggested that non-repeat coverage may be noisy for some samples, and we thus plotted the coverage of reads from all strains around and over the Undecim cluster of wPauO11 (Figure S9). Since the coverage was approximately doubled compared to the surrounding single-copy regions for all *w*Pau strain variants, the same genomic region is likely duplicated in all sequenced *w*Pau genomes.

## Discussion

Neotropical *Drosophila* species of the willistoni and saltans groups were previously shown to carry closely related *Wolbachia* strains with high similarity to the *w*Au strain from *D. simulans* (Miller and Riegler 2006). Using 14 new *Wolbachia* genomes from five Neotropical *Drosophila* species, we confirm that these strains are highly similar and belong to a monophyletic clade including strain *w*Au – i.e., the *w*Au-likes.

### The *w*Au-like *Wolbachia* show patterns of co-speciation and horizontal transmission

To investigate the possibility of co-speciation between the *w*Au-like *Wolbachia* strains infecting willistoni and saltans group flies, *w*Au-like_ws_ strains, and their hosts, we inferred a phylogeny and compared it to the nuclear and mitochondrial phylogenies of the hosts (Baião et al. 2023). We found that the phylogeny of *w*Au-like_ws_ strains was congruent with the nuclear phylogeny of the host species, except for the *Wolbachia* strain *w*Pau and its host *D. paulistorum*. Complete congruence was, however, found between the topologies of the two maternally transmitted genetic entities, *Wolbachia* and mitochondria. Hence, based on phylogenetic congruence, it seems plausible that a *w*Au-like strain was present in the ancestor of the willistoni and saltans groups at their divergence ca. 20 million years ago (Powell et al. 2003; Suvorov et al. 2022) and that this strain has co-diverged with the current hosts ever since. However, phylogenetic congruence can be achieved in the absence of co- speciation, a phenomenon referred to as pseudo-co-speciation and is especially common if host switching preferentially occurs between closely related species (De Vienne et al. 2007). Thus, to infer true co-speciation, temporal congruence between the two compared entities should be observed. However, since no reliable estimate of the mutation rate exists for any *Wolbachia* strain, dating the nodes in our tree will be highly unreliable. Nevertheless, to attempt to investigate temporal congruence, we calculated the synonymous substitution rate of *w*Au-like_ws_ strains relative to that of their hosts and compared that to previously reported cases of *Wolbachia*-host co-speciation. We found the relative rates in our dataset to be quite constant between different *Wolbachia*-host pairs, with *D. paulistorum- D. willistoni* and their *Wolbachia* having the highest ratio, around 3-7 times higher than the others. Even so, all rates were between one and two orders of magnitude lower than in previous examples of co- speciation (Raychoudhury et al. 2008; Gerth and Bleidorn 2016; Conner et al. 2017). Thus, if all suggested co-speciation examples were true, the *w*Au-like_ws_ strains either have evolved slower than other co-speciating *Wolbachia* strains or the willistoni and saltans group flies have evolved faster than estimated in previous studies. However, we don’t know if the earlier reported cases represent true examples of co-speciation, since neither has proven temporal congruence. Additionally, mutation and substitution rates as well as generation times can vary widely between different organisms, even between very closely related bacterial and animal species. Hence, we also calculated mutation rates of *Wolbachia,* based on the synonymous substitution rates and the substitution rate in single-copy genes, assuming that *Wolbachia* divergence times are congruent with host divergence times. Using synonymous substitutions, we got mutation rates between 10^-10^–10^-8^ substitutions per site per year and using all substitutions in single-copy genes, we got even lower estimates of the mutation rate of 3-7x10^-11^ substitutions per site per year. Both rates are lower than the estimated mutation rate in other bacteria (Gibson and Eyre-Walker 2019) as well as in *Wolbachia* strain *w*Mel, where the mutation rate was estimated from population data to be between 2.88x10^-9^ –

1.29x10^-8^ substitutions per site per year (Richardson et al. 2012). However, Richardson et al., note that the *w*Mel rate is biased, as it includes slightly deleterious mutations only observed at the population level, and not at the species level. Consequently, for our calculated rates to match the generational mutation rates of bacteria (ranging between 10^-10^– 10^-8^ synonymous substitutions per synonymous site per generation (Lynch et al. 2023), all *Wolbachia* strains analysed here, except the ones in *Nasonia,* must have a generation time of almost a year. Since this is even longer than the generation times of the hosts, our estimated mutation rates are likely incorrect. There are indeed many potential sources of error for our mutation rate estimates, for example, synonymous substitutions could be under strong selection, or the recombination rate could be extremely high and eliminate diversity between strains, both of which could yield underestimated mutation rates. Additionally, the host divergence times could be wrong, which, depending on whether they are over- or under-estimated, would result in either lower or higher mutation rates. At this point, we can’t completely rule out any of these factors, although selection seems unlikely given that the effective population size of *Wolbachia* is probably low, and it is hard to understand how recombination between the *w*Au-like_ws_ strains could be so rampant if our *Wolbachia* strains are only vertically transmitted. However, the divergence times of the hosts are likely a smaller problem, as they would have to be off by orders of magnitude for our estimated rates to be comparable to the rates of *w*Mel and other bacteria. Alternatively, for co-speciation to be believable, most of the *Wolbachia* strains analysed here must have a very low mutation rate or long generation time, or a combination thereof. If not, neither the *w*Au-like_ws_ strains nor the *Wolbachia* strains from *Nomada* and *Nasonia* have co-speciated with their hosts but may instead have been horizontally transmitted into their hosts via introgression or alternative mechanisms, as suggested in other similar systems (Turelli et al. 2018; Cooper et al. 2019).

It is unlikely that *Wolbachia* could have been recently transmitted between the relatively distantly related species of the willistoni and saltans group via introgression, but the topological congruence between mitochondria and *Wolbachia* could suggest that introgressions have facilitated at least some horizontal transmission events. As previously reported, *D. paulistorum* has two mitotypes, α and β, which were probably both introgressed from an unknown species within the willistoni group (Baião et al. 2023). Hence, it is possible that *w*Pau was introgressed into *D. paulistorum* from a willistoni group species together with one of the mitotypes. Potentially, the β mitotype, since it is most common among *D. paulistorum* semispecies and all *D. paulistorum* lines in this study carry it. As *w*Pau has complete and thus likely functional *cif* genes, the spread of *w*Pau and the replacement of the ancestral *D. paulistorum* mitochondrion could potentially have been driven by CI. However, to fully answer this question, data from additional *D. paulistorum* semispecies, especially the ones carrying the α-mitotype, are needed.

Additionally, at least four horizontal transmission events of *w*Au-like *Wolbachia*, i.e., *w*Inn, *w*Bor, *w*Inc, and *w*Au, between distantly related *Drosophila* species have likely occurred relatively recently, as previously suggested (Ballard 2004; Miller and Riegler 2006; Sheeley and McAllister 2009; Wallau et al. 2016), through mechanisms other than introgression. We can only speculate about how these *w*Au-like *Wolbachia* strains infected and spread in these hosts, but CI is not a likely mechanism for the spread since none of them carry what appear to be functional *cif* genes. The well-studied strain *w*Au has long been known to not induce CI in its native host, *D. simulans* (Hoffmann et al. 1996). However, it does provide high antiviral protection in its host (Martinez et al., 2015), a potentially beneficial phenotype that might have facilitated its spread. Nothing is currently known about the phenotypic effects *w*Inc has on its host, but both *w*Inn and *w*Bor cause male-killing, which could have contributed to their spread. Furthermore, *w*Inn was also seen to protect the host against RNA viruses (Unckless and Jaenike 2012). It is also possible that *w*Au-like strains provide yet unknown beneficial host effects.

In conclusion, more research is required to determine whether *Wolbachia* switches hosts so frequently that pseudo-co-speciation is observed independently in multiple clades and to identify mechanisms that facilitate the transfer and spread of *Wolbachia* infections.

### The genome of the obligate *Wolbachia* strain *w*Pau is expanded and has some genomic features similar to recently host-associated symbionts

When free-living bacteria become host-associated, many of their genes become superfluous in the new nutrient-rich, stable environment of the host and are hence lost or pseudogenized due to lack of selection (McCutcheon and Moran 2012). In addition, when symbionts transition from free-living to host-associated, they experience a reduction in effective population size that increases the effect of genetic drift (Moran and Plague 2004, Wernegreen 2015). Consequently, their genomes tend to accumulate slightly deleterious mutations, including many IS element insertions (Bobay and Ochman 2017). However, over time, often after becoming obligate for their hosts, symbiont genomes generally shrink further in size and lose repeated sequences and pseudogenes (McCutcheon and Moran 2012, Bobay and Ochman 2017), leading to a conserved gene order. This stage of the genome reduction process can, for example, be observed in obligate mutualistic *Wolbachia* of nematodes that have small genomes (Werren et al. 2008, Dudzic et al. 2022) with few repeats and mobile elements (Lefoulon et al. 2020) compared to most facultative *Wolbachia* strains.

To our knowledge, *w*Pau is the only *Wolbachia* strain found in the willistoni and saltans group of flies that is obligate for the host (Miller et al. 2010). Hence, *w*Pau is expected to have a smaller genome than its close facultative relatives. Contrary to the expectation, we found that *w*PauO11 has the largest genome among the *w*Au-like *Wolbachia* strains with complete genomes. Genome size increase in *Wolbachia* is generally associated with a higher proportion of prophage WO (Kampfraath et al. 2019; Lefoulon et al. 2020; Wolfe et al. 2021; Vancaester and Blaxter 2023), but in *w*Pau, we rather found a reduction of prophage WO content. Instead, we attribute the increase in genome size in *w*Pau to a still ongoing expansion of an IS4 element that makes up almost 10% of the genome and has greatly affected the gene order. The extensive IS4 element proliferation, high rate of genomic rearrangements and reduced coding percentage in the *w*Pau genome are genomic features that are more common in recently host-associated facultative symbionts than evolutionary persistent obligate symbionts. All of them, however, suggest a large influence of genetic drift, potentially caused by a reduction in effective population size. We can think of at least three possible reasons why the effective population size of *w*Pau could have decreased compared to its close relatives.

Firstly, under the assumption of horizontal transmission, *w*Pau might have reduced the effective population size since only a few *Wolbachia* cells likely established the infection in *D. paulistorum* initially. While we don’t see an expansion of IS elements in other *w*Au-like strains with complete genomes that have likely shifted host recently, i.e., *w*Inn and *w*Au, the *w*Inn genome shares other features with the *w*Pau genome, including many rearrangements and low coding density. Possibly, *w*Inn had more repeats but lost them over time due to deletions. Although these similarities with *w*Inn might indicate that horizontal transmission could initially cause a higher effect of genetic drift, we do not believe this to be the main reason for the sustained high levels of genetic drift in *w*Pau.

Secondly, the transition from facultative to obligate host association could have yielded a reduction in the effective population size of *w*Pau. Several similar genome features can be found in the genome of the mutualistic *Wolbachia* strain *w*Cle infecting bedbugs (Hosokawa et al. 2010), including an expansion of IS elements (with over 100 copies of IS5) and reduced prophage WO content (Lefoulon et al. 2020). Potential IS element expansions are also recorded in other obligate symbiont genomes, for example, the co-obligate *Serratia symbiotica* strain Ct from the aphid *Cinara tujafilina* (Manzano-Marín and Latorre 2014), where several IS element families have expanded (Renoz et al. 2021) and some species of the genus *Sodalis* (Oakeson et al. 2014; Koga and Moran 2014; Ankrah et al. 2018; Manzano-Marín et al. 2018; Garber et al. 2021). However, since no genome sequences of closely related facultative strains are available in any of these cases, the IS element expansion can’t be linked to a transition to obligate host association. Thirdly, a reduction in the effective population size of *w*Pau could be a consequence of a reduction in the effective population size of its host, which might be linked to the fact that *D. paulistorum* is a superspecies consisting of multiple fully or partly reproductively isolated semispecies. Currently, no data about the effective population sizes of any species in the willistoni or saltans groups exist; however, indirect evidence indicates a lower effective population size in *D. paulistorum* compared to other willistoni and saltans group species. Large eukaryotic genomes with a high proportion of transposable elements often occur in species with low effective population sizes (Lynch and Conery 2003), due to the high influence of genetic drift that allows mobile elements to accumulate. Since *D. paulistorum* has the largest genome with the highest repetitive content of the sequenced willistoni group species (Kim et al. 2021, Gebert et al. 2024), we might speculate that *D. paulistorum* has a lower effective population size than other willistoni and saltans group species. If so, this could explain not only the high rate of IS4 element insertions before the divergence of the *w*Pau strain variants but also the continued high rate that seems to be ongoing in *w*Pau even after its likely introgression and transition from facultative to obligate.

### Prophage WO genes may be important for the obligate symbiosis of *w*Pau

The WO prophage is known to play an important role in facultative *Wolbachia* host associations, given that it carries genes directly responsible for reproductive manipulation (Beckmann et al. 2017; LePage et al. 2017; Perlmutter et al. 2019; Fricke and Lindsey 2024), infection titer and virulence (Chrostek and Teixeira 2015; Duarte et al. 2021). For the obligate *w*Pau, we demonstrate loss of prophage WO genes compared to closely related facultative strains. Specifically, *w*Pau has lost almost an entire copy (WOa) and some parts of the other (WOb), suggesting a general lack of selection on WO-associated genes rather than any specific WO prophage region. In contrast, nine of the eleven genes of the prophage WO-associated Undecim cluster are instead duplicated in *w*PauO11 as well as the other *w*Pau strain variants. Although the functions of the Undecim cluster genes are currently unknown, they are thought to play a role in *Wolbachia*-host interactions, as they, among other functions, encode proteins potentially involved in exopolysaccharide and/or lipopolysaccharide biosynthesis (Bordenstein and Bordenstein 2022). Previous work has also shown that the Undecim cluster has been horizontally transferred between several unrelated insect endosymbionts and that the genes are constitutively highly expressed in *Wolbachia* strain *w*Mel during the whole life cycle of *D. melanogaster* (Gutzwiller et al. 2015). The fact that this prophage WO-associated set of genes is duplicated in *w*Pau, while many other prophage WO genes are deleted, suggests that the region could encode proteins potentially important for the obligate symbiosis between *w*Pau and its host. If true, prophage WO might thus contribute genes for the persistence of a *Wolbachia* infection, even in the absence of CI or other reproductive manipulations.

## Conclusions

We conclude that the ancestor of Neotropical *Drosophila* in the willistoni and saltans groups may have been infected with a *w*Au-like *Wolbachia* and co-speciated with their hosts, except for the *w*Pau strain in *D. paulistorum*, which was likely introgression from another species within the willistoni group together with one of the two mitotypes. However, such an interpretation suggests that *Wolbachia* must have a very low mutation rate. Since the real mutation rate of any strain of *Wolbachia i*s still unknown, further research is needed to determine if co-speciation is a possible interpretation.

We also found that contrary to expectations, the genome of the obligate *w*Pau strain was larger than its facultative relatives due to the ongoing proliferation of an IS4 element, typically associated with genomes of recently host-associated symbionts. A reduced coding percentage and more pseudogenes are also hallmarks of the early stages of genome reduction, all of which are likely due to a reduction in the effective population size.

In general, the effective population size can be reduced multiple times after a bacterium becomes host associated. Thus, the genome reduction process might take place in a punctuated manner, as episodes of effective population size reduction can temporarily lead to an increased genome size via repeat expansion, and a decrease in the proportion of coding content, only to be reversed by homologous recombination at such repeats over time. Consequently, only when mobile elements and repeats are lost from a genome altogether can this cycle be broken.

## Data availability

All *Wolbachia* genomes and annotations have been submitted to a public sequence database and will be made available upon publication. The mitochondrial genomes and nuclear data are available in NCBI Bioproject PRJNA643793 (Baiaõ et al. 2023). The accession numbers of other *Wolbachia* genomes used in our analyses can be found in Supplementary Tables S3 and S4.

## Abbreviations

ACT: Artemis Comparison Tool
ANKs: ankyrin repeat domain-containing genes
CI: Cytoplasmic Incompatibility
EAM: Eukaryotic Association Module
ONT: Oxford Nanopore Technologies
SNP: Single Nucleotide Polymorphism
MLST: Multi-Locus Strain-Typing
gCF: Gene Concordance Factor
sCF: Site Concordance Factor
*w*Au-like_ws_: *Wolbachia* strains infecting fruit flies of the willistoni and saltans groups
WOa/b: WO Phage sequences orthologous to WOMelA and WOMelB, respectively

## Declarations

### Ethics approval and consent to participate

Not applicable for this study.

## Consent for publication

Not applicable for this study.

### Competing Interests

The authors declare that they have no competing interests.

## Funding

This work was supported by the Swedish research council VR grant 2014-4353 to LK and by the Austrian Science Fund FWF grant P28255B22 to WJM.

## Acknowledgements

We thank Mercè Montoliu Nerín for assembling the *Drosophila paulistorum* O11 genome from which the *w*PauO11 genome contig was extracted and for helpful discussions throughout the project, Tom Martin for performing the DNA extraction for Nanopore sequencing of *D. paulistorum* O11, Lina Juzokaite for *Wolbachia*-enriched DNA extractions for PacBio sequencing, and Illumina sequencing. We also thank Athanasios Alexiou for assembling the first version of the *w*WilP98 genome. Sequencing was performed by the SNP&SEQ Technology Platform and Uppsala Genome Center (UGC) in Uppsala, Sweden. The facilities are part of the National Genomics Infrastructure (NGI) Sweden and Science for Life Laboratory. Some of the data handling was enabled by resources provided by the Swedish National Infrastructure for Computing (SNIC) at UPPMAX. SNP&SEQ, UGC and Uppmax are supported by the Swedish Research Council and SNP&SEQ are supported by the Knut and Alice Wallenberg Foundation.

## Author contribution

LK and WJM conceived and designed the study. KP and LK performed assemblies. KP and LK performed annotations. KP and LK performed comparative analyses. KP performed phy- logenetic analyses. LK and KP wrote the paper with contributions from WJM. All authors read and approved the manuscript.

## Notes

### Competing Interest Statement

The authors have declared no competing interest.

### Summary of Updates

Mutation rate calculations using a Bayesian method calibrated with the host divergence times were added. We replaced the Wolbachia strain tree in Figure 2 with one containing only the willistoni and saltans group Wolbachia strains. We added a table that contains the divergence times and the mutation rates. We made some minor textual adjustments throughout the manuscript.

## References

Baião GC, Janice J, Galinou M, Klasson L. Comparative Genomics Reveals Factors Associated with Phenotypic Expression of Wolbachia. Genome Biology and Evolution. 2021 July 1;13(7):evab111.

Baião GC, Schneider DI, Miller WJ, Klasson L. The effect of Wolbachia on gene expression in Drosophila paulistorum and its implications for symbiont-induced host speciation. BMC Genomics. 2019 Dec;20(1):465.

Baião GC, Schneider DI, Miller WJ, Klasson L. Multiple introgressions shape mitochondrial evolutionary history in Drosophila paulistorum and the Drosophila willistoni group. Molecular Phylogenetics and Evolution. 2023 Mar;180:107683.

Bakovic V, Schebeck M, Stauffer C, Schuler H. Wolbachia-Mitochondrial DNA Associations in Transitional Populations of Rhagoletis cerasi. Insects. Multidisciplinary Digital Publishing Institute; 2020 Oct;11(10):675.

Baldo L, Dunning Hotopp JC, Jolley KA, Bordenstein SR, Biber SA, Choudhury RR, et al. Multilocus Sequence Typing System for the Endosymbiont Wolbachia pipientis. Applied and Environmental Microbiology. American Society for Microbiology; 2006 Nov;72(11):7098–110.

Ballard JWO. Sequential Evolution of a Symbiont Inferred From the Host: Wolbachia and Drosophila simulans. Molecular Biology and Evolution. 2004 Mar 1;21(3):428–42.

Balvín O, Roth S, Talbot B, Reinhardt K. Co-speciation in bedbug Wolbachia parallel the pattern in nematode hosts. Sci Rep. 2018 June 11;8(1):8797.

Bankevich A, Nurk S, Antipov D, Gurevich AA, Dvorkin M, Kulikov AS, et al. SPAdes: a new genome assembly algorithm and its applications to single-cell sequencing. J Comput Biol. 2012 May;19(5):455–77.

Bateman A, Birney E, Cerruti L, Durbin R, Etwiller L, Eddy SR, et al. The Pfam protein families database. Nucleic Acids Res. 2002 Jan 1;30(1):276–80.

Beckmann JF, Ronau JA, Hochstrasser M. A Wolbachia Deubiquitylating Enzyme Induces Cytoplasmic Incompatibility. Nat Microbiol. 2017 Mar 1;2:17007.

Blasco-Costa I, Hayward A, Poulin R, Balbuena JA. Next-generation cophylogeny: unravelling eco-evolutionary processes. Trends in Ecology & Evolution. 2021 Oct 1;36(10):907–18.

Bobay L-M, Ochman H. The Evolution of Bacterial Genome Architecture. Front. Genet. [Internet]. 2017 [cited 2023 Aug 9];8. Available from: https://www.frontiersin.org/articles/10.3389/fgene.2017.00072

Bordenstein SR, Bordenstein SR. Eukaryotic association module in phage WO genomes from Wolbachia. Nat Commun. 2016 Oct 11;7(1):13155.

Bordenstein SR, Bordenstein SR. Widespread phages of endosymbionts: Phage WO genomics and the proposed taxonomic classification of Symbioviridae. Matic I, editor. PLoS Genet. 2022 june 6;18(6):e1010227.

Bourtzis K, Nirgianaki A, Markakis G, Savakis C. Wolbachia infection and cytoplasmic incompatibility in Drosophila species. Genetics. 1996 Nov;144(3):1063–73.

Bruen TC, Philippe H, Bryant D. A simple and robust statistical test for detecting the presence of recombination. Genetics. 2006 Apr;172(4):2665–81.

Camacho C, Coulouris G, Avagyan V, Ma N, Papadopoulos J, Bealer K, et al. BLAST+: architecture and applications. BMC Bioinformatics. 2009 Dec 15;10(1):421.

Cariou M, Duret L, Charlat S. The global impact of Wolbachia on mitochondrial diversity and evolution. Journal of Evolutionary Biology. 2017;30(12):2204–10.

Carver TJ, Rutherford KM, Berriman M, Rajandream M-A, Barrell BG, Parkhill J. ACT: the Artemis Comparison Tool. Bioinformatics. 2005 Aug 15;21(16):3422–3.

Charlat S, Duplouy A, Hornett EA, Dyson EA, Davies N, Roderick GK, et al. The joint evolutionary histories of Wolbachia and mitochondria in Hypolimnas bolina. BMC Evolutionary Biology. 2009 Mar 24;9(1):64.

Charleston MA. Cospeciation. Encyclopedia of Evolutionary Biology [Internet]. Elsevier; 2016 [cited 2023 Sept 25]. p. 381–6. Available from: https://linkinghub.elsevier.com/retrieve/pii/B9780128000496002006

Chrostek E, Marialva MSP, Yamada R, O’Neill SL, Teixeira L. High Anti-Viral Protection without Immune Upregulation after Interspecies Wolbachia Transfer. PLOS ONE. Public Library of Science; 2014 June 9;9(6):e99025.

Chrostek E, Teixeira L. Mutualism Breakdown by Amplification of Wolbachia Genes. Malik HS, editor. PLoS Biol. 2015 Feb 10;13(2):e1002065.

Conner WR, Blaxter ML, Anfora G, Ometto L, Rota-Stabelli O, Turelli M. Genome comparisons indicate recent transfer of Ri-like Wolbachia between sister species Drosophila suzukii and D. subpulchrella. Ecology and Evolution. John Wiley & Sons, Ltd; 2017 Nov;7(22):9391–404.

Cooper BS, Vanderpool D, Conner WR, Matute DR, Turelli M. Wolbachia Acquisition by Drosophila yakuba-Clade Hosts and Transfer of Incompatibility Loci Between Distantly Related Wolbachia. Genetics. 2019 Aug;212(4):1399–419.

Danecek P, Bonfield JK, Liddle J, Marshall J, Ohan V, Pollard MO, et al. Twelve years of SAMtools and BCFtools. GigaScience. 2021 Jan 29;10(2):giab008.

De Vienne DM, Giraud T, Shykoff JA. When can host shifts produce congruent host and parasite phylogenies? A simulation approach. Journal of Evolutionary Biology. 2007;20(4):1428–38.

Drillon G, Carbone A, Fischer G. SynChro: A Fast and Easy Tool to Reconstruct and Visualize Synteny Blocks along Eukaryotic Chromosomes. PLOS ONE. Public Library of Science; 2014 Mar 20;9(3):e92621.

Drillon G, Champeimont R, Oteri F, Fischer G, Carbone A. Phylogenetic Reconstruction Based on Synteny Block and Gene Adjacencies. Mol Biol Evol. 2020 Sept 1;37(9):2747–62.

Duarte EH, Carvalho A, López-Madrigal S, Costa J, Teixeira L. Forward genetics in Wolbachia: Regulation of Wolbachia proliferation by the amplification and deletion of an addictive genomic island. PLOS Genetics. Public Library of Science; 2021 June 18;17(6):e1009612.

Dudzic JP, Curtis CI, Gowen BE, Perlman SJ. A highly divergent *Wolbachia* with a tiny genome in an insect-parasitic tylenchid nematode. Proc. R. Soc. B. 2022 Sept 28;289(1983):20221518.

Ellegaard KM, Klasson L, Näslund K, Bourtzis K, Andersson SGE. Comparative Genomics of Wolbachia and the Bacterial Species Concept. Matic I, editor. PLoS Genet. 2013 Apr 4;9(4):e1003381.

Emms DM, Kelly S. OrthoFinder: phylogenetic orthology inference for comparative genomics. Genome Biology. 2019 Nov 14;20(1):238.

Fisher RM, Henry LM, Cornwallis CK, Kiers ET, West SA. The evolution of host-symbiont dependence. Nat Commun. 2017 July 4;8:15973.

Frank AC, Amiri H, Andersson SGE. Genome deterioration: loss of repeated sequences and accumulation of junk DNA. Genetica. 2002;

Fricke LC, Lindsey ARI. Identification of Parthenogenesis-Inducing Effector Proteins in Wolbachia. Genome Biology and Evolution. 2024 Apr 1;16(4):evae036.

Gavotte L, Henri H, Stouthamer R, Charif D, Charlat S, Boulétreau M, et al. A Survey of the Bacteriophage WO in the Endosymbiotic Bacteria Wolbachia. Molecular Biology and Evolution. 2007 Feb 1;24(2):427–35.

Gerth M, Bleidorn C. Comparative genomics provides a timeframe for Wolbachia evolution and exposes a recent biotin synthesis operon transfer. Nat Microbiol. 2016 Dec 22;2(3):16241.

Gibson B, Eyre-Walker A. Investigating Evolutionary Rate Variation in Bacteria. J Mol Evol. 2019 Dec 1;87(9):317–26.

Gordon D, Abajian C, Green P. Consed: a graphical tool for sequence finishing. Genome Res. 1998 Mar;8(3):195–202.

Gutzwiller F, Carmo CR, Miller DE, Rice DW, Newton ILG, Hawley RS, et al. Dynamics of Wolbachia pipientis Gene Expression Across the Drosophila melanogaster Life Cycle. G3 (Bethesda). 2015 Oct 23;5(12):2843–56.

Guy L, Roat Kultima J, Andersson SGE. genoPlotR: comparative gene and genome visualization in R. Bioinformatics. 2010 Sept 15;26(18):2334–5.

Hill T, Unckless RL, Perlmutter JI. Positive Selection and Horizontal Gene Transfer in the Genome of a Male-Killing Wolbachia. Molecular Biology and Evolution. 2022 Jan 1;39(1):msab303.

Hosokawa T, Koga R, Kikuchi Y, Meng X-Y, Fukatsu T. Wolbachia as a bacteriocyte- associated nutritional mutualist. Proceedings of the National Academy of Sciences. Proceedings of the National Academy of Sciences; 2010 Jan 12;107(2):769–74.

Huerta-Cepas J, Serra F, Bork P. ETE 3: Reconstruction, Analysis, and Visualization of Phylogenomic Data. Molecular Biology and Evolution. 2016 June 1;33(6):1635–8.

Ilinsky Y. Coevolution of Drosophila melanogaster mtDNA and Wolbachia Genotypes. PLOS ONE. Public Library of Science; 2013 Jan 17;8(1):e54373.

Kampfraath AA, Klasson L, Anvar SY, Vossen RHAM, Roelofs D, Kraaijeveld K, et al. Genome expansion of an obligate parthenogenesis-associated Wolbachia poses an exception to the symbiont reduction model. BMC Genomics. 2019 Dec;20(1):106.

Katoh K, Toh H. Recent developments in the MAFFT multiple sequence alignment program. Briefings in Bioinformatics. 2008 July 1;9(4):286–98.

Kaur R, Shropshire JD, Cross KL, Leigh B, Mansueto AJ, Stewart V, et al. Living in the endosymbiotic world of Wolbachia: A centennial review. Cell Host & Microbe. 2021 June;29(6):879–93.

Kim BY, Wang JR, Miller DE, Barmina O, Delaney E, Thompson A, et al. Highly contiguous assemblies of 101 drosophilid genomes. Coop G, Wittkopp PJ, Sackton TB, editors. eLife. eLife Sciences Publications, Ltd; 2021 July 19;10:e66405.

Lefoulon E, Bain O, Makepeace BL, d’Haese C, Uni S, Martin C, et al. Breakdown of coevolution between symbiotic bacteria Wolbachia and their filarial hosts. PeerJ. PeerJ Inc.; 2016 Mar 28;4:e1840.

Lefoulon E, Clark T, Guerrero R, Cañizales I, Cardenas-Callirgos JM, Junker K, et al. Diminutive, degraded but dissimilar: Wolbachia genomes from filarial nematodes do not conform to a single paradigm. Microb Genom. 2020 Dec;6(12):mgen000487.

LePage DP, Metcalf JA, Bordenstein SR, On J, Perlmutter JI, Shropshire JD, et al. Prophage WO genes recapitulate and enhance Wolbachia-induced cytoplasmic incompatibility. Nature. Nature Publishing Group; 2017 Mar;543(7644):243–7.

Li H, Durbin R. Fast and accurate short read alignment with Burrows-Wheeler transform. Bioinformatics. 2009 July 15;25(14):1754–60.

Lynch M, Ali F, Lin T, Wang Y, Ni J, Long H. The divergence of mutation rates and spectra across the Tree of Life. EMBO reports. John Wiley & Sons, Ltd; 2023 Oct 9;24(10):e57561.

Lynch M, Conery JS. The origins of genome complexity. Science. 2003 Nov 21;302(5649):1401–4.

Maechler M, Rousseeuw P, Struyf A, Hubert M, Hornik K. cluster: Cluster Analysis Basics and Extensions [Internet]. 2021. Available from: https://CRAN.R-project.org/package=cluster

Manzano-Marín A, D’acier AC, Clamens A-L, Orvain C, Cruaud C, Barbe V, et al. A Freeloader? The Highly Eroded Yet Large Genome of the Serratia symbiotica Symbiont of Cinara strobi. Genome Biology and Evolution. Oxford University Press; 2018 Sept;10(9):2178.

Manzano-Marín A, Latorre A. Settling Down: The Genome of Serratia symbiotica from the Aphid Cinara tujafilina Zooms in on the Process of Accommodation to a Cooperative Intracellular Life. Genome Biology and Evolution. 2014 July 1;6(7):1683–98.

Marçais G, Delcher AL, Phillippy AM, Coston R, Salzberg SL, Zimin A. MUMmer4: A fast and versatile genome alignment system. PLOS Computational Biology. Public Library of Science; 2018 Jan 26;14(1):e1005944.

Martinez J, Ant TH, Murdochy SM, Tong L, Filipe A da S, Sinkins SP. Genome sequencing and comparative analysis of Wolbachia strain wAlbA reveals Wolbachia-associated plasmids are common. PLOS Genetics. Public Library of Science; 2022 Sept 19;18(9):e1010406.

Martinez J, Klasson L, Welch JJ, Jiggins FM. Life and Death of Selfish Genes: Comparative Genomics Reveals the Dynamic Evolution of Cytoplasmic Incompatibility. Molecular Biology and Evolution. 2021 Jan 4;38(1):2–15.

Martinez J, Ok S, Smith S, Snoeck K, Day JP, Jiggins FM. Should Symbionts Be Nice or Selfish? Antiviral Effects of Wolbachia Are Costly but Reproductive Parasitism Is Not. McGraw EA, editor. PLoS Pathog. 2015 July 1;11(7):e1005021.

McCutcheon JP, Moran NA. Extreme genome reduction in symbiotic bacteria. Nat Rev Microbiol. 2012 Jan;10(1):13–26.

Miller WJ, Ehrman L, Schneider D. Infectious Speciation Revisited: Impact of Symbiont- Depletion on Female Fitness and Mating Behavior of Drosophila paulistorum. Parrish C, editor. PLoS Pathog. 2010 Dec 2;6(12):e1001214.

Miller WJ, Riegler M. Evolutionary Dynamics of w Au-Like Wolbachia Variants in Neotropical Drosophila spp. Appl Environ Microbiol. 2006 Jan;72(1):826–35.

Minh BQ, Schmidt HA, Chernomor O, Schrempf D, Woodhams MD, Von Haeseler A, et al. IQ-TREE 2: New Models and Efficient Methods for Phylogenetic Inference in the Genomic Era. Teeling E, editor. Molecular Biology and Evolution. 2020 May 1;37(5):1530–4.

Moran NA, Plague GR. Genomic changes following host restriction in bacteria. Current Opinion in Genetics & Development. 2004 Dec;14(6):627–33.

Müller MJ, von Mühlen C, Valiati VH, Valente VL da S. Wolbachia pipientis is associated with different mitochondrial haplotypes in natural populations of Drosophila willistoni. Journal of Invertebrate Pathology. 2012 Jan 1;109(1):152–5.

Nikoh N, Hosokawa T, Moriyama M, Oshima K, Hattori M, Fukatsu T. Evolutionary origin of insect–Wolbachia nutritional mutualism. Proceedings of the National Academy of Sciences. Proceedings of the National Academy of Sciences; 2014 July 15;111(28):10257–62.

Perlmutter JI, Bordenstein SR, Unckless RL, LePage DP, Metcalf JA, Hill T, et al. The phage gene wmk is a candidate for male killing by a bacterial endosymbiont. PLOS Pathogens. Public Library of Science; 2019 Sept 10;15(9):e1007936.

Powell JR, Sezzi E, Moriyama EN, Gleason JM, Caccone A. Analysis of a Shift in Codon Usage in Drosophila. J Mol Evol. 2003 Aug 1;57(1):S214–25.

R Core Team. R: A Language and Environment for Statistical Computing [Internet]. Vienna, Austria: R Foundation for Statistical Computing; 2018. Available from: https://www.R-project.org/

Rambaut A. FigTree v1.4: Tree Figure Drawing Tool. [Internet]. 2009. Available from: http://tree.bio.ed.ac.uk/software/figtree/

Raychoudhury R, Baldo L, Oliveira DCSG, Werren JH. Modes of Acquisition of *Wolbachia*: Horizontal Transfer, Hybrid Introgression, and Codivergence in the *Nasonia* Species Complex. Evolution. 2008 Sept;63(1):165–83.

Renoz F, Arai H, Pons I. The genus Sodalis as a resource for understanding the multifaceted evolution of bacterial symbiosis in insects. Symbiosis. 2024 Mar 1;92(2):187– 208.

Renoz F, Foray V, Ambroise J, Baa-Puyoulet P, Bearzatto B, Mendez GL, et al. At the Gate of Mutualism: Identification of Genomic Traits Predisposing to Insect-Bacterial Symbiosis in Pathogenic Strains of the Aphid Symbiont Serratia symbiotica. Front. Cell. Infect. Microbiol. [Internet]. Frontiers; 2021 June 29 [cited 2024 Aug 8];11. Available from: https://www.frontiersin.org/journals/cellular-and-infectionmicrobiology/articles/10.3389/fcimb.2021.660007/full

Richardson MF, Weinert LA, Welch JJ, Linheiro RS, Magwire MM, Jiggins FM, et al. Population Genomics of the Wolbachia Endosymbiont in Drosophila melanogaster. PLOS Genetics. Public Library of Science; 2012 Dec 20;8(12):e1003129.

Russell SL, Chappell L, Sullivan W. A symbiont’s guide to the germline. Current Topics in Developmental Biology [Internet]. Elsevier; 2019 [cited 2023 July 18]. p. 315–51. Available from: https://linkinghub.elsevier.com/retrieve/pii/S0070215319300559

Ryan JA, Ulrich JM. quantmod: Quantitative Financial Modelling Framework [Internet]. 2022. Available from: https://CRAN.R-project.org/package=quantmod

Schneider DI, Ehrman L, Engl T, Kaltenpoth M, Hua-Van A, Le Rouzic A, et al. Symbiont- Driven Male Mating Success in the Neotropical Drosophila paulistorum Superspecies. Behav Genet. 2019 Jan;49(1):83–98.

Seemann T. Prokka: rapid prokaryotic genome annotation. Bioinformatics. 2014 July 15;30(14):2068–9.

Sheeley SL, McAllister BF. Mobile male-killer: similar Wolbachia strains kill males of divergent Drosophila hosts. Heredity. Nature Publishing Group; 2009 Mar;102(3):286–92.

Shen X-X, Hittinger CT, Rokas A. Contentious relationships in phylogenomic studies can be driven by a handful of genes. Nat Ecol Evol. Nature Publishing Group; 2017 Apr 10;1(5):1– 10.

Shropshire JD, Leigh B, Bordenstein SR. Symbiont-mediated cytoplasmic incompatibility: What have we learned in 50 years? eLife. 2020;9:e61989.

Small ST, Tisch DJ, Zimmerman PA. Molecular epidemiology, phylogeny and evolution of the filarial nematode Wuchereria bancrofti. Infect Genet Evol. 2014 Dec;0:33–43.

Suvorov A, Kim BY, Wang J, Armstrong EE, Peede D, D’Agostino ERR, et al. Widespread introgression across a phylogeny of 155 *Drosophila* genomes. Current Biology. 2022 Jan 10;32(1):111–123.e5.

Syberg-Olsen MJ, Garber AI, Keeling PJ, McCutcheon JP, Husnik F. Pseudofinder: Detection of Pseudogenes in Prokaryotic Genomes. Molecular Biology and Evolution. 2022 July 1;39(7):msac153.

Turelli M, Cooper BS, Richardson KM, Ginsberg PS, Peckenpaugh B, Antelope CX, et al. Rapid Global Spread of *w*Ri-like *Wolbachia* across Multiple *Drosophila*. Current Biology. 2018 Mar 19;28(6):963–971.e8.

Turelli M, Hoffmann AA. Cytoplasmic incompatibility in Drosophila simulans: dynamics and parameter estimates from natural populations. Genetics. 1995 Aug 1;140(4):1319–38.

Unckless RL, Boelio LM, Herren JK, Jaenike J. Wolbachia as populations within individual insects: causes and consequences of density variation in natural populations. Proceedings of the Royal Society B: Biological Sciences. Royal Society; 2009 May 6;276(1668):2805–11.

Unckless RL, Jaenike J. Maintenance of a Male-Killing Wolbachia in Drosophila Innubila by Male-Killing Dependent and Male-Killing Independent Mechanisms. Evolution. 2012 Mar 1;66(3):678–89.

Vancaester E, Blaxter M. Phylogenomic analysis of Wolbachia genomes from the Darwin Tree of Life biodiversity genomics project. Teixeira L, editor. PLoS Biol. 2023 Jan 23;21(1):e3001972.

Wallau GL, da Rosa MT, De Ré FC, Loreto ELS. Wolbachia from Drosophila incompta: just a hitchhiker shared by Drosophila in the New and Old World? Insect Molecular Biology. 2016;25(4):487–99.

Waterhouse AM, Procter JB, Martin DMA, Clamp M, Barton GJ. Jalview Version 2—a multiple sequence alignment editor and analysis workbench. Bioinformatics. 2009 May 1;25(9):1189–91.

Wernegreen JJ. Endosymbiont evolution: predictions from theory and surprises from genomes: Endosymbiont genome evolution. Ann. N.Y. Acad. Sci. 2015 Dec;1360(1):16–35.

Werren JH, Baldo L, Clark ME. Wolbachia: master manipulators of invertebrate biology. Nat Rev Microbiol. 2008 Oct;6(10):741–51.

Wickham H, Averick M, Bryan J, Chang W, McGowan LD, François R, et al. Welcome to the tidyverse. Journal of Open Source Software. 2019;4(43):1686.

Wolfe TM, Bruzzese DJ, Klasson L, Corretto E, Lečić S, Stauffer C, et al. Comparative genome sequencing reveals insights into the dynamics of Wolbachia in native and invasive cherry fruit flies. Molecular Ecology. 2021;30(23):6259–72.

Wu M, Sun LV, Vamathevan J, Riegler M, Deboy R, Brownlie JC, et al. Phylogenomics of the Reproductive Parasite Wolbachia pipientis wMel: A Streamlined Genome Overrun by Mobile Genetic Elements. PLOS Biology. Public Library of Science; 2004 Mar 16;2(3):e69.

Xie Z, Tang H. ISEScan: automated identification of insertion sequence elements in prokaryotic genomes. Bioinformatics. 2017 Nov 1;33(21):3340–7.

Yang Z. PAML 4: Phylogenetic Analysis by Maximum Likelihood. Molecular Biology and Evolution. 2007 Apr 18;24(8):1586–91.

Zeng Y, Wiens JJ. Do mutualistic interactions last longer than antagonistic interactions? Proceedings of the Royal Society B: Biological Sciences. Royal Society; 2021 Sept 8;288(1958):20211457.

Zhang Y-K, Ding X-L, Zhang K-J, Hong X-Y. Wolbachia Play an Important Role in Affecting mtDna Variation of Tetranychus truncatus (Trombidiformes: Tetranychidae). Environmental Entomology. 2013 Dec 1;42(6):1240–5.

