## Supplementary material for "A co-speciation dilemma and a lifestyle transition with genomic consequences in *Wolbachia* of Neotropical *Drosophila*": Figure S

### Supplemental figures

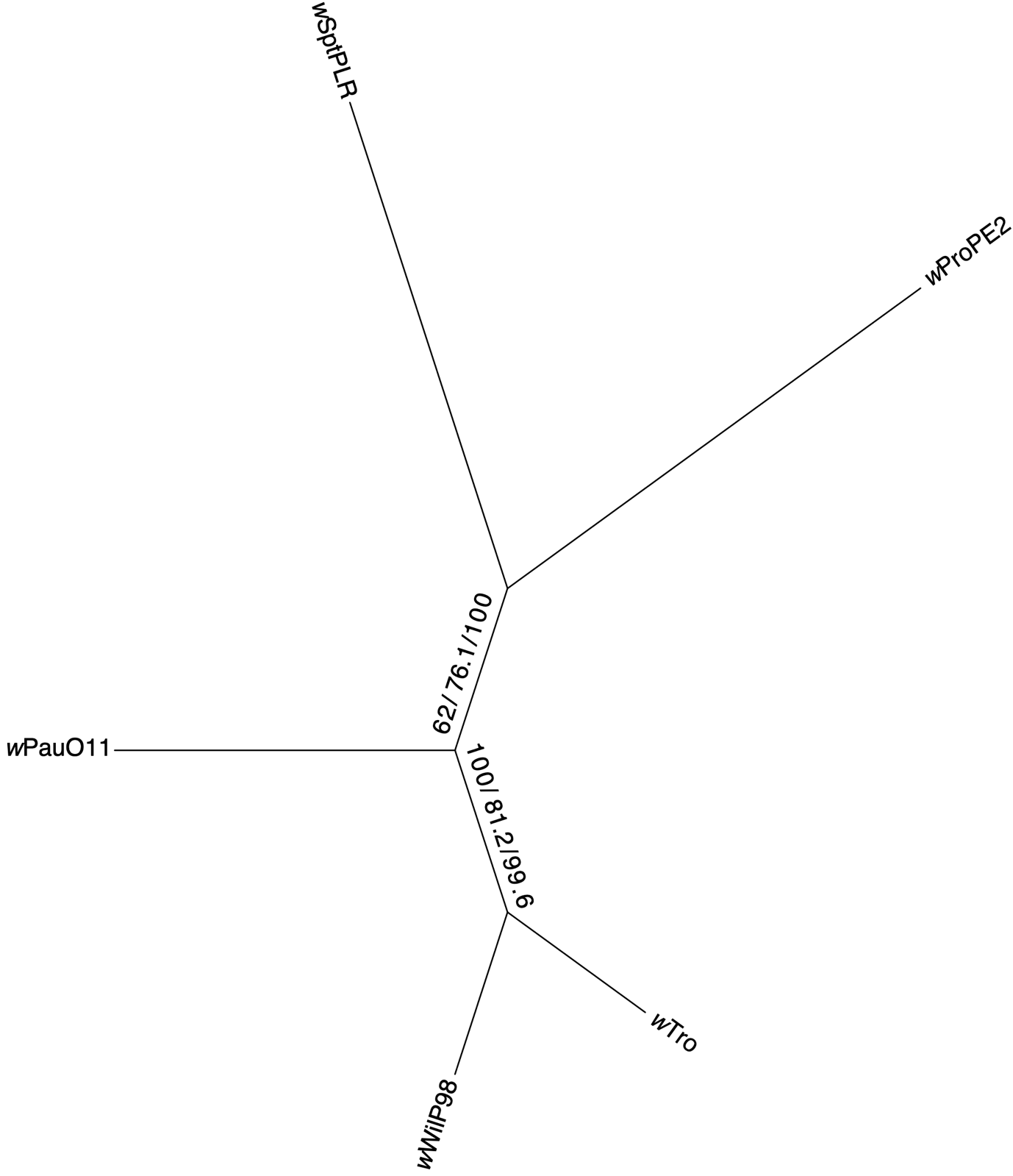

**Figure S1. Gene and site concordance factors for the *w*Au-like strains.** A subtree of the main phylogeny in Figure 1, including only *w*Au-like_ws_ strains. Numbers on the branches are the bootstrap, gCF and sCF values.

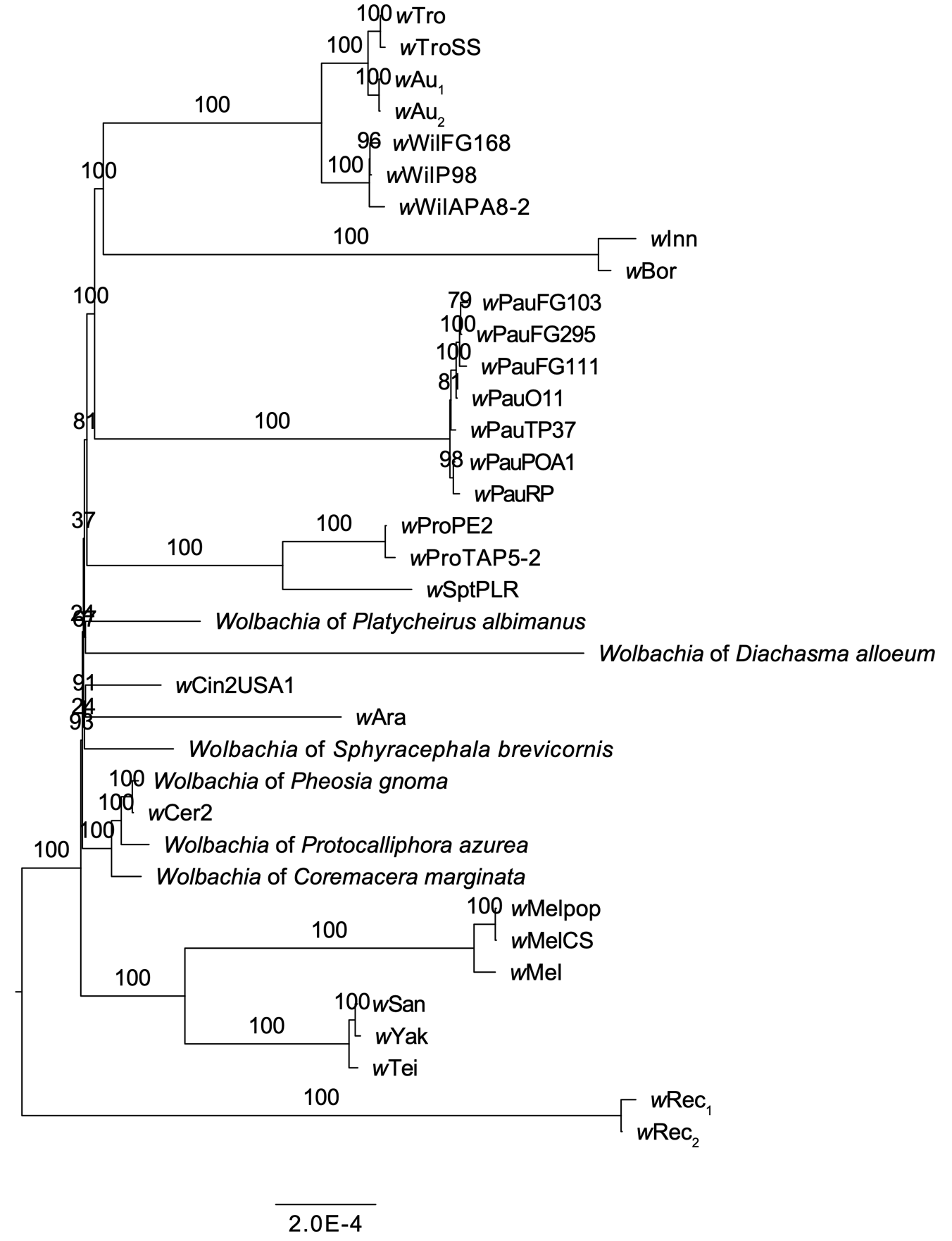

**Figure S2. Maximum-likelihood phylogeny of *w*Au-like strains (without *w*Inc) and related *Wolbachia*.** The tree was inferred from an alignment of 620 single-copy orthologs with IQ-TREE (same as in Figure 1 but excluding the strain *w*Inc). Support values from 1000 ultrafast bootstrap replicates are displayed on the nodes.

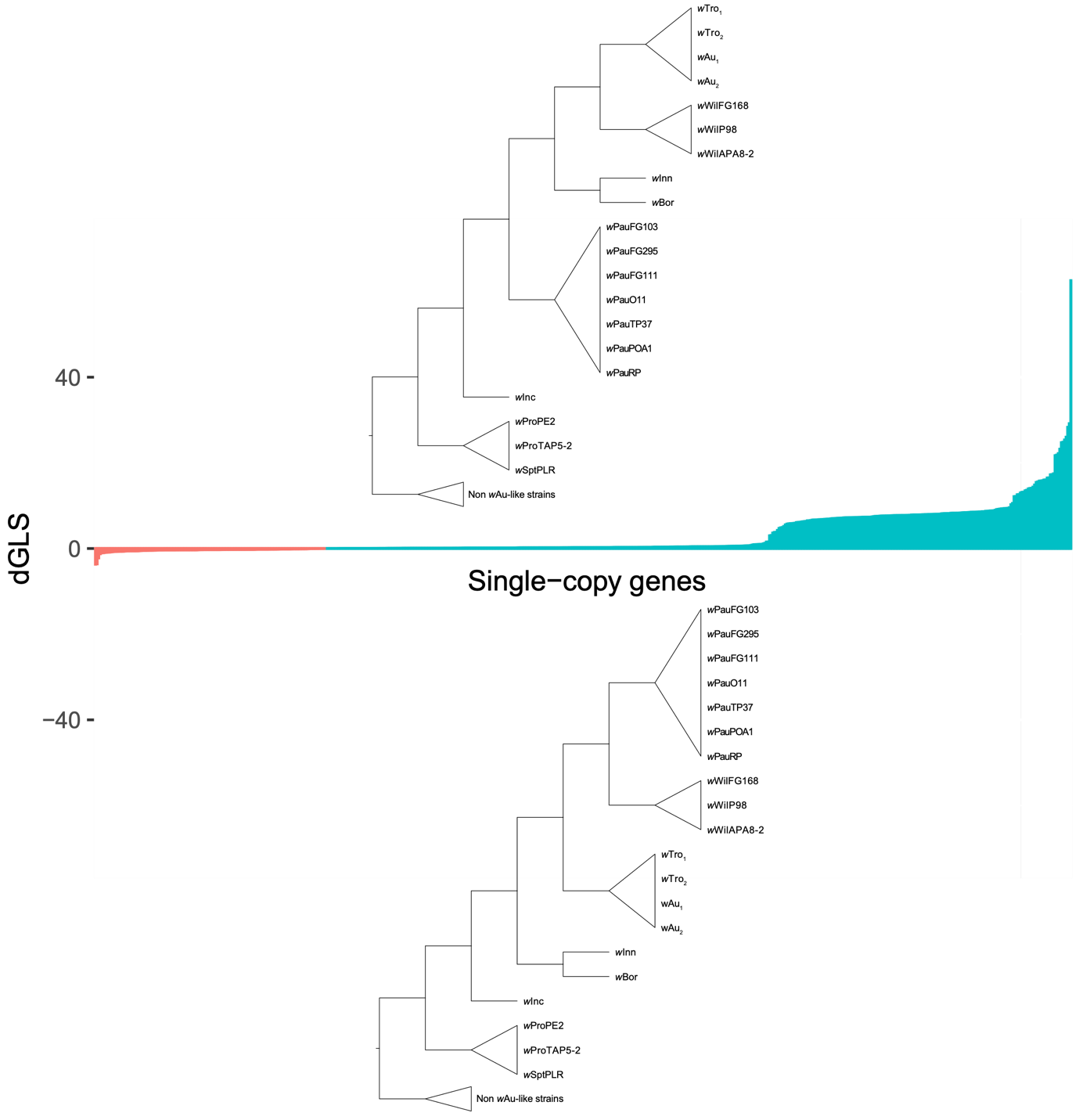

**Figure S3. Gene-wise log-likelihood scores for *w*Au-like topology.** For each of the 620 single-copy genes used to infer the concatenated *w*Au-like strain phylogeny, we plot the difference in likelihood between two test topologies, one that looks like the inferred *Wolbachia* strain topology (in blue) and an alternative topology that follows the host nuclear phylogeny (in red). Scores for all 620 single-copy genes are sorted in ascending order along the X-axis. Most genes support the inferred topology for the *Wolbachia* *w*Au-like strains (top tree) over the host nuclear topology (bottom tree).

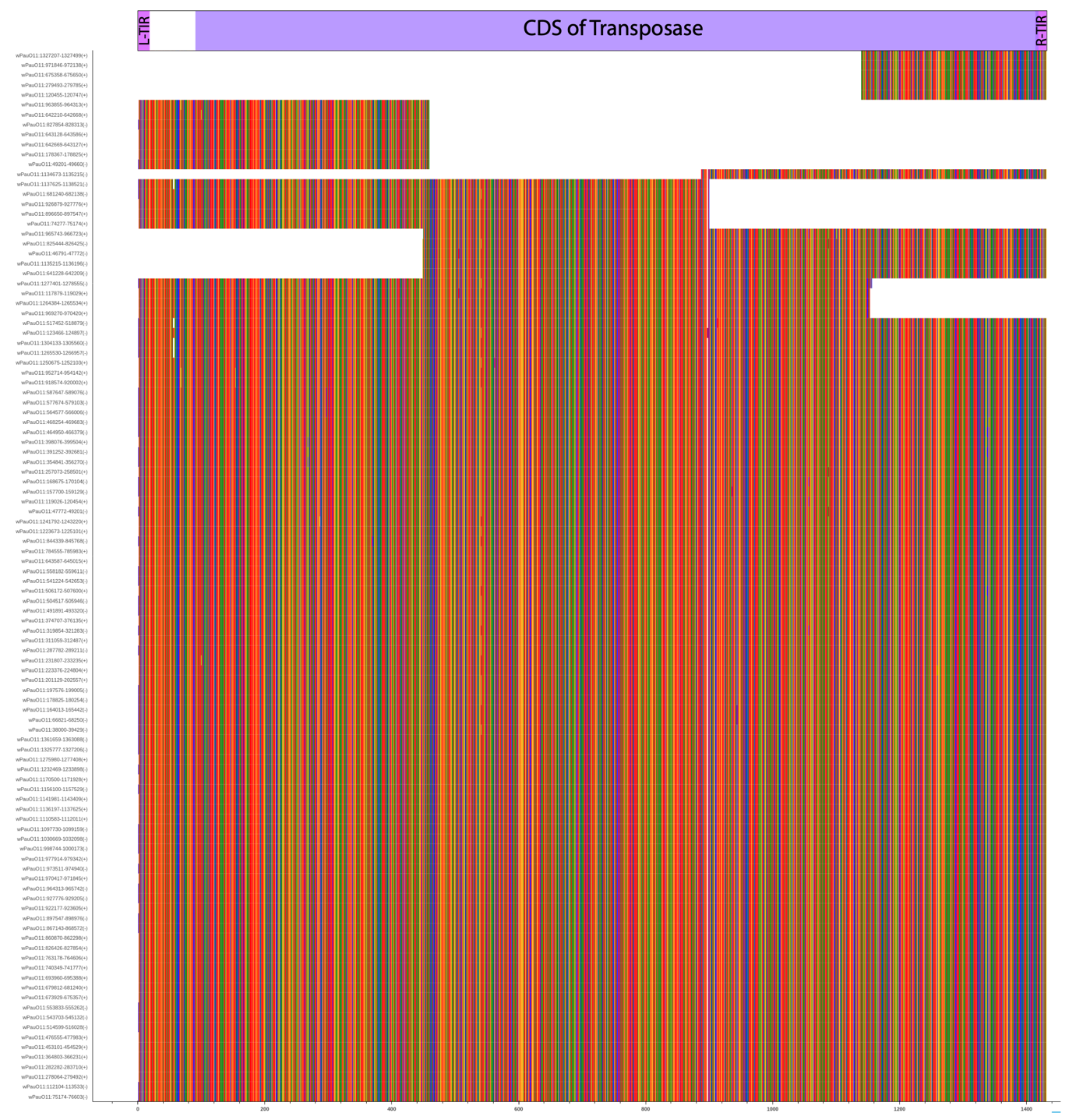

**Figure S4. Nucleotide alignment of all IS4 elements identified in *w*PauO11.** Alignment of 106 nucleotide sequences that ISEscan identified as complete or partial IS4 elements. On top of the alignment, we show the left and right Terminal Inverted Repeats (TIRs) as well as the coding sequence (CDS) of the transposase for the IS4 element. For each element, the position in the genome is reported. By matching incomplete copies, we estimate the number of complete IS4 elements to be 96.

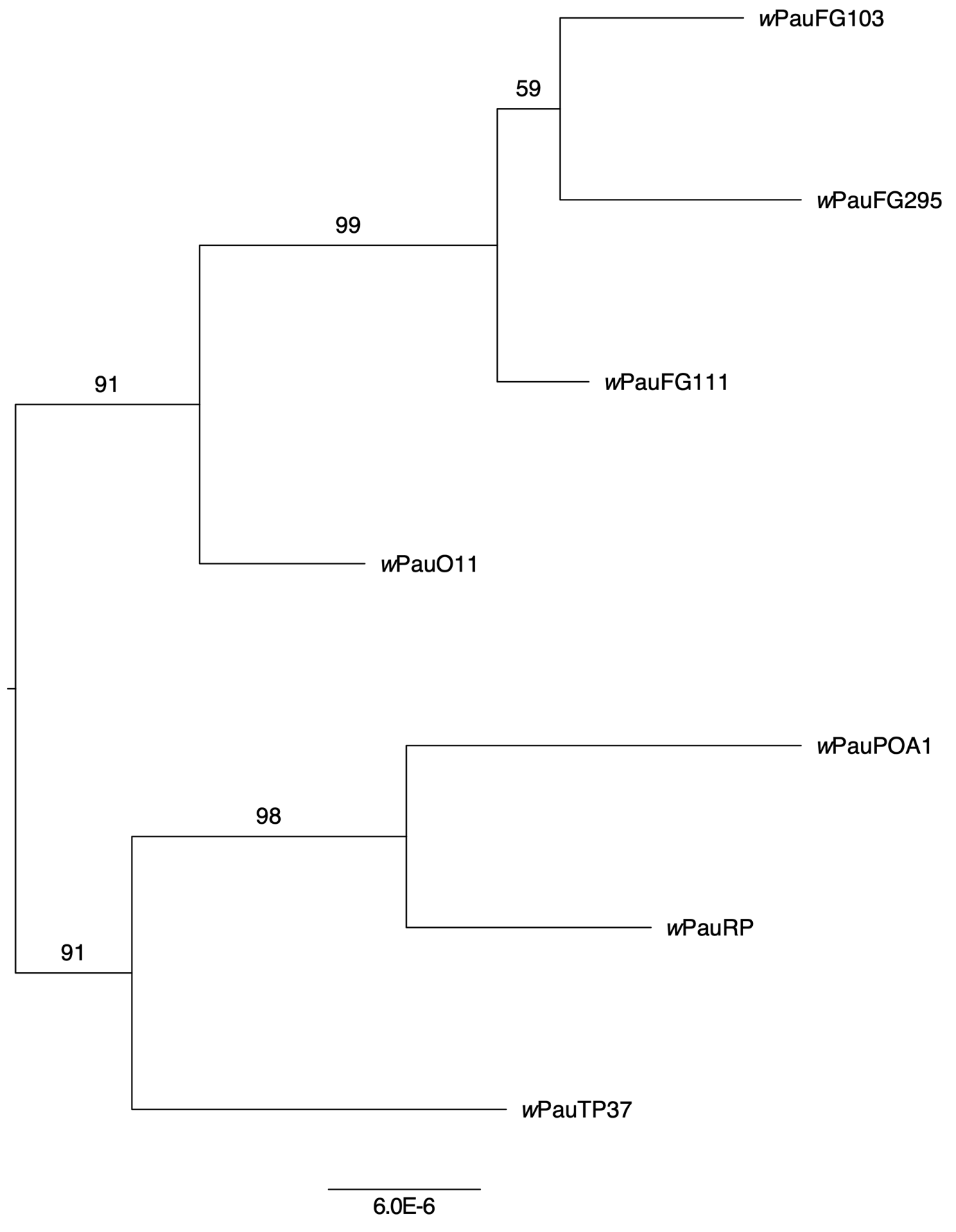

**Figure S5. A phylogeny of the *w*Pau strains based on 93 genome-wide SNPs.** IQTREE was used to produce the tree and support values from 1000 ultrafast bootstrap replicates are displayed on the nodes. Midpoint rooting is applied.

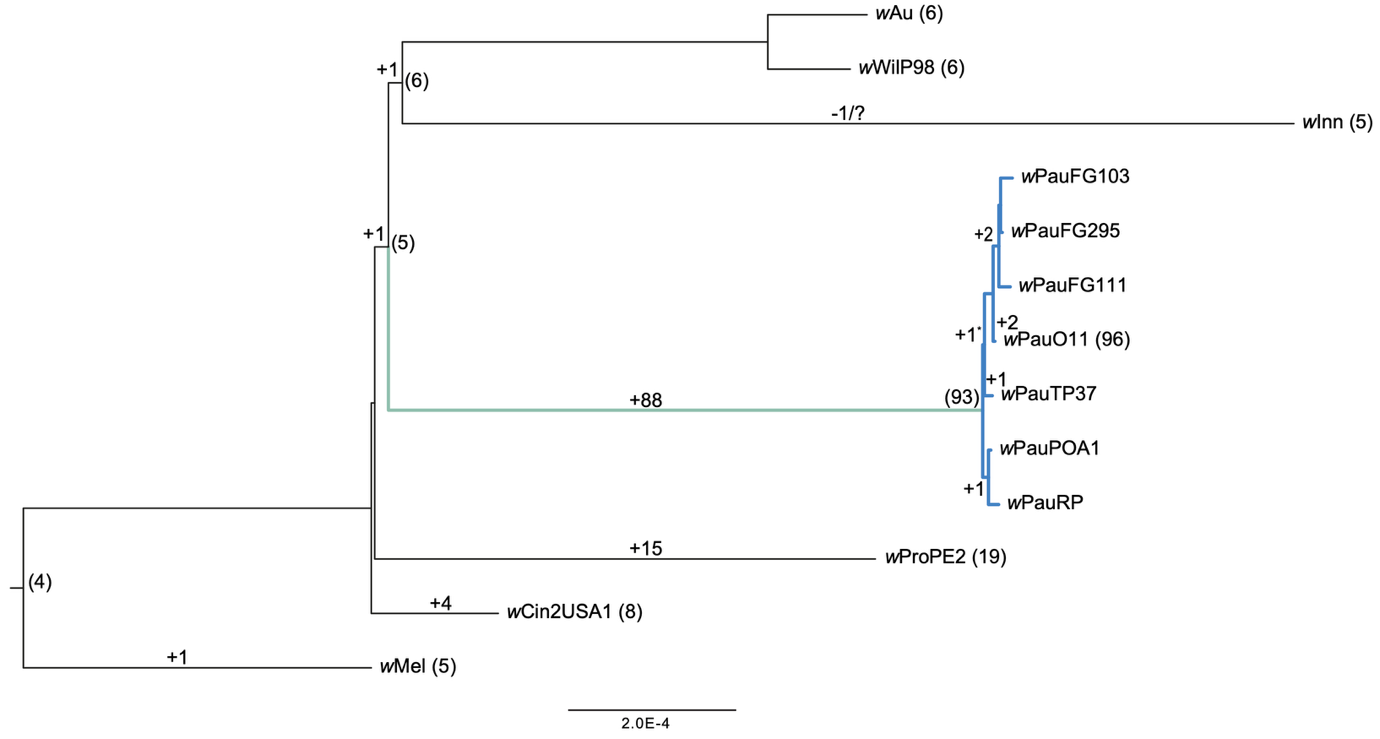

**Figure S6. Reconstruction of IS4 insertion events across the phylogeny of the *w*Au-like strains.** Numbers in parentheses at the nodes show the inferred number of IS4 elements in ancestral genomes. Numbers on the branches show inferred IS4 element insertion events for each lineage. Numbers in parentheses on branch tips show the number of IS4 elements detected for each genome by ISEscan.

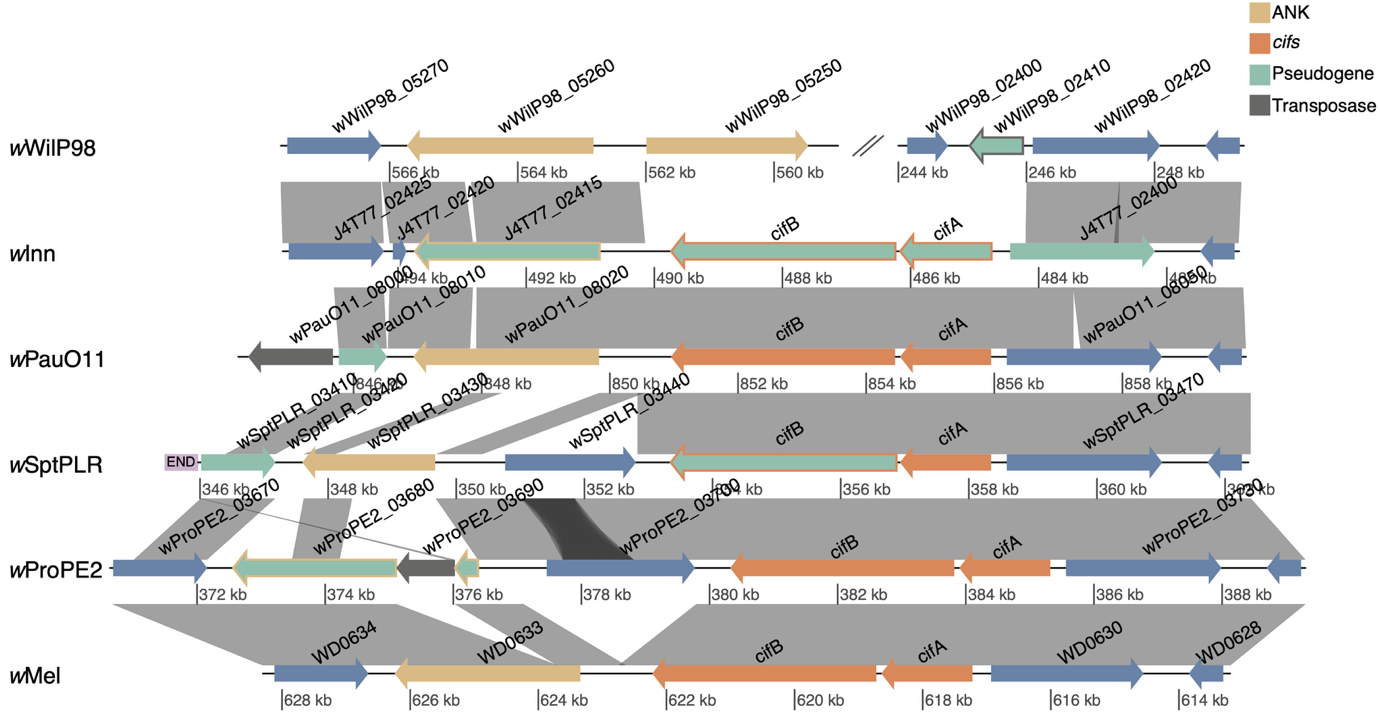

**Figure S7. The *cifA* and *cifB* genes in the genomes of *w*Au-likes and *w*Mel.** Genes are depicted with arrows, the cif genes in orange, ankyrin-domain containing genes in yellow, transposases in grey, and other protein-coding genes in blue. Genes in green are pseudogenes. Similarities from blastn searches are shown with grey ribbons between the genomes. The figure was made with genoPlotR (Guy et al. 2010).

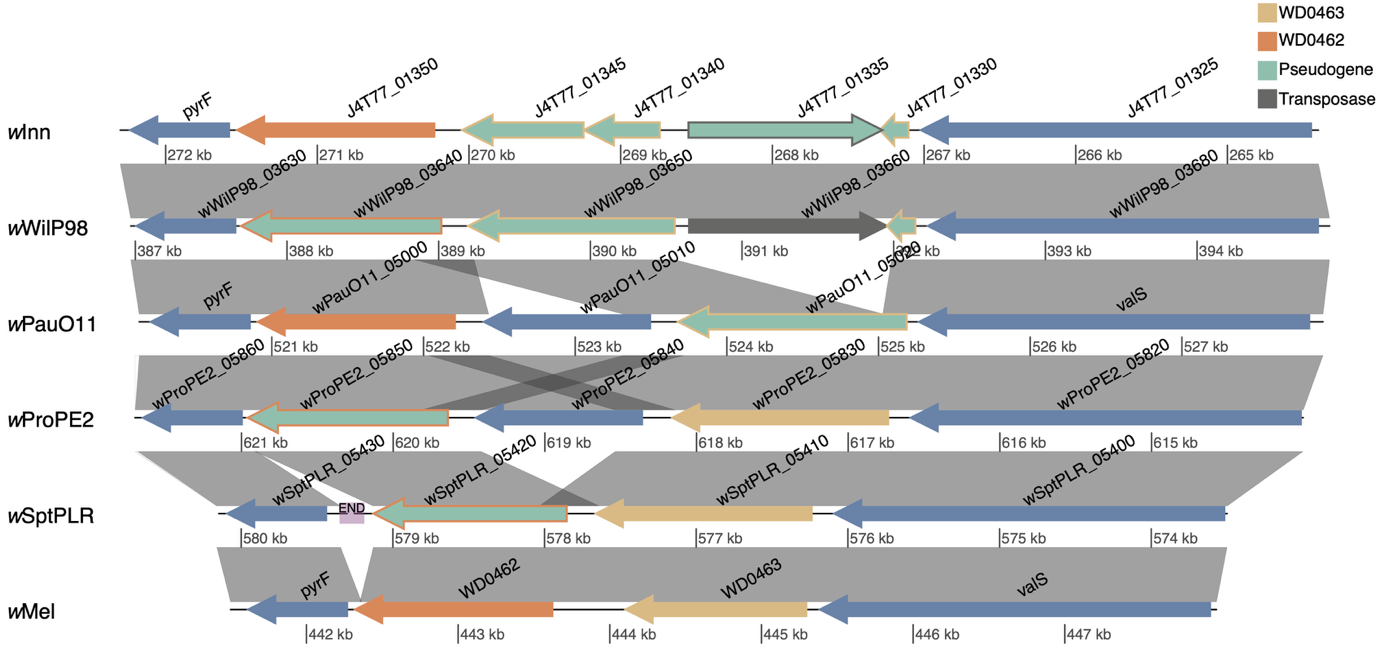

**Figure S8. The WD0462 and WD0463 genes in genomes of *w*Au-likes and *w*Mel.** Genes are depicted with arrows, homologues to WD0462 in orange, homologues to WD0463 in yellow, transposases in grey, and other protein-coding genes in blue. Genes in green are pseudogenes. Similarities from blastn searches are shown with grey ribbons between the genomes. The figure was made with genoPlotR (Guy et al. 2010).

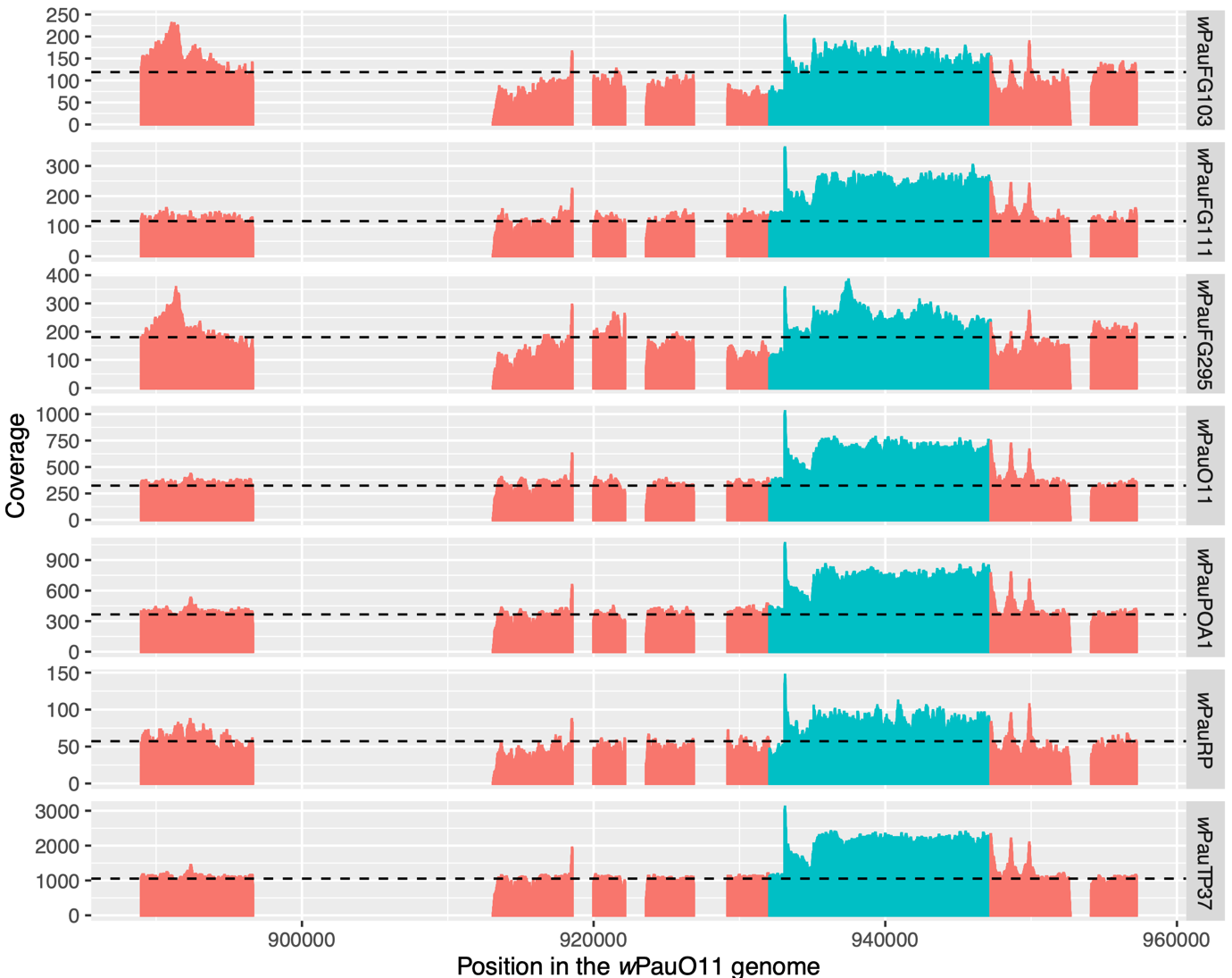

**Figure S9. Read coverage of *w*Pau strain variants mapped on the *w*PauO11 genome.** All IS4 elements and the first copy of the Undecim cluster were masked in wPauO11. The coverage over the unmasked Undecim cluster copy is shown in blue, and the coverage over other parts of the genome is shown in red. The dashed line shows the average read coverage for non-repeated regions of the genome.
